# Autoimmune inflammation triggers aberrant astrocytic calcium signaling to impair synaptic plasticity

**DOI:** 10.1101/2023.08.01.551248

**Authors:** AM Baraibar, T Colomer, A Moreno-García, A Bernal-Chico, E Sánchez, C Utrilla, R Serrat, E Soria-Gómez, A Rodríguez-Antigüedad, A Araque, C Matute, G Marsicano, S Mato

**Author notes:** These authors contributed equally: Andrés M Baraibar, Teresa Colomer. Correspondence (G.M.) (S.M.).

## Abstract

Cortical pathology involving inflammatory and neurodegenerative mechanisms is a hallmark of multiple sclerosis (MS) and a correlate of disease progression and cognitive decline. Astrocytes play a pivotal role in MS initiation and progression but astrocyte-neuronal network alterations contributing to gray matter pathology remain undefined. Here we measured astrocytic calcium in the experimental autoimmune encephalomyelitis (EAE) model of MS using fiber photometry in freely behaving mice and two-photon imaging *ex vivo*. We identified the emergence of spontaneously hyperactive cortical astrocytes displaying calcium transients of increased duration as well as dysfunctional responses to cannabinoid, glutamate and purinoreceptor agonists during acute EAE disease. Deficits in astrocyte calcium responses are associated to abnormal signaling by G_i_ and G_q_ protein coupled receptors in the inflamed cortex and are partially mirrored in cells activated with pro-inflammatory factors both *in vitro* and *ex vivo* thus suggesting cell-autonomous effects of the cortical neuroinflammatory environment. Finally, we show that deregulated astrocyte calcium activity is associated to an enhancement of glutamatergic gliotransmission and a shift of astrocyte-mediated short-term and long-term plasticity mechanisms towards synaptic potentiation. Overall our data identities astrocyte-neuronal network dysfunction as key pathological feature of the inflammatory gray matter that may contribute to MS symptomatology and clinical progression.

## Introduction

Multiple sclerosis (MS) is a chronic demyelinating disease of the central nervous system (CNS) initiated by pathogenic immune cell responses against myelin followed by a broader inflammatory and neurodegenerative process (Mahad et al., 2015). Over the last decades a number of neuroimaging and histopathological studies have demonstrated that cortical pathology is centrally involved in MS symptomatology and progression (Calabrese et al., 2010; De Stefano et al., 2003). Functional and structural alterations affecting the cortex are present from the earliest stages of MS, contribute to motor, sensory and cognitive deficits and predict the accumulation of disability (De Stefano et al., 2003; Eshaghi et al., 2018; Feinstein et al., 2014). Neocortical gray matter atrophy in MS involves intermingled inflammatory and neurodegenerative mechanisms that include early excitatory-inhibitory imbalance and widespread synapse loss as key pathological features (Calabrese et al., 2015; Ellwardt et al., 2018; Jafari et al., 2021; Jürgens et al., 2016; Potter et al., 2016). However, the molecular, cellular and circuit mechanisms that drive dysregulated cortical network activity in MS remain unclear.

Astrocytes support synaptic function and plasticity within neuronal circuits in a broad range of physiological settings. It has long been recognized that astroglial cells respond to synaptically released neurotransmitters with intracellular calcium elevations that culminate in the release of neuroactive substances such as glutamate, adenosine/ATP and D-serine (Araque et al., 2014). The release of these factors termed gliotransmitters enables astrocyte calcium signaling to modulate synaptic strength and fine-tune neuronal network activity in a tightly controlled manner. Astrocytes undergo a pronounced transformation in the context of neuroinflammation whereby they adopt a reactive phenotypes and exhibit important disease-promoting functions that contribute to neurological disability (Linnerbauer et al., 2020; Sofroniew, 2020). Research on the role of reactive astrocytes in MS has established that these cells drive inflammatory lesion formation and promote the neurodegenerative process by multiple mechanisms that include neurotoxicity and dysfunctional astrocyte-to-neuron crosstalk (Chao et al., 2019; Pitt et al., 2000; Ponath et al., 2018). Nevertheless, current understanding on how aberrant interactions between astrocytes and neuronal cells contribute to MS symptomatology and disease progression remains poor and deficits in astrocyte-mediated gliotransmitter release and modulation of synaptic function remain underexplored.

Recent advances in functional calcium imaging have led to novel insights how aberrant astrocytic activity patterns may contribute to neuronal functional deficits in the diseased CNS (Shigetomi et al., 2016). In this study we interrogated the deficits in astrocyte calcium dynamics and astrocyte-to-neuron synaptic crosstalk associated to cortical pathology in MS using a combination of imaging modalities applied to the experimental autoimmune encephalomyelitis (EAE) model of the disease. We show that autoimmune inflammation markedly alters the intrinsic calcium activity of cortical astrocytes that displayed spontaneous oscillations of increase duration but reduced responses to neurotransmitter receptor agonists at acute EAE disease. Our results also show that these alterations emerge in response to the inflammatory environment and rely on G protein mediated calcium signaling defects. Finally, we show that dysfunctional astrocyte calcium dynamics encompass aberrant glutamate gliotransmission and deregulation astrocyte-to-neuron plasticity mechanisms towards the potentiation of synaptic excitation. Overall, these data unveil cortical astrocyte-neuron network dysregulation in MS and suggest a novel mechanism of gray matter pathology relying on aberrant astrocyte mediated modulation of synaptic function during autoimmune inflammation.

## Methods

### Mice

All experiments were performed in accordance with the Guide for the Care and Use of Laboratory Animals (National Research Council Committee, 2011) and the European Communities Council Directive of September 22th 2010 (2010/63/EU74). Experiments were approved by the local ethical committees of the University of the Basque Country (approval numbers 2017140, M202017144 and 2022245) and the University of Bordeaux (approval number A33063098) and the French Ministry of Agriculture and Forestry (authorization number 3306369). Female naive mice on a C57BL/6N background (Janvier, France), inositol 1,4,5-trisphosphate (IP_3_) receptor type 2 knockout (IP_3_R_2_-KO) mice (generously donated by Dr. Gertrudis Perea; Instituto Cajal, Madrid, Spain), constitutive CB_1_-KO and CB_1_-WT mice, CB_1_^f/f^ mice (carrying the ‘‘floxed’’ *Cnr1* gene) and inducible GFAP-CB_1_-KO mutant mice and GFAP-CB_1_-WT littermates (7-14 weeks of age) were used. Cages were enriched and mice were maintained under standard conditions (food and water *ad libitum*; 12 h-12 h light-dark cycle). Experiments were performed during dark cycle (light off at 8:00 h a.m.). Pups for primary cell cultures (P4-P5) were obtained from homozygote CB_1_-WT pairs. The number of mice in each experimental group was similar. No statistical methods were used to predetermine sample size.

GFAP-CB_1_-KO mice were generated by crossing CB ^f/f^ mice (Marsicano et al., 2002) with GFAP-CreERT2 mice (Hirrlinger et al., 2006) using a three-step backcrossing procedure to obtain CB_1_^f/f;GFAP-CreERT2^ and CB ^f/f^ littermates, called GFAP-CB_1_-KO and GFAP-CB_1_-WT. Deletion of the *Cnr1* gene was obtained in adult mice (8-9 weeks of age) by daily intraperitoneal (i.p.) injections of tamoxifen (1 mg dissolved at 10 mg/ml in 90% sesame oil, 10% ethanol) for 7 days (Han et al., 2012; Robin et al., 2018).

### Drugs

THC was obtained from THC Pharm GmbH (Frankfurt, Germany). ATP disodium salt-hydrate (ATP), thapsigargin and WIN55-212-2 (WIN) were purchased from Sigma-Aldrich (St-Louis, USA); tetrodotoxin (TTX), AM251, clozapine-N-oxide (CNO), (S)DHPG, glutamate monosodium salt-hydrate and LY354740 from Tocris (Bristol, UK). For *in vitro* and *ex vivo* experiments, WIN, AM251 and CNO were dissolved in DMSO; TTX was dissolved in ddH_2_O, ATP and LY354740 and glutamate were dissolved in recording media. The concentration of DMSO was never higher than 0.001%. For *in vivo* administration, THC was prepared freshly before the experiments and was dissolved in a mixture of 5% ethanol, 4% cremophor and saline. Doses and concentrations of the different drugs were chosen based on previous published data or preliminary experiments.

### EAE model

Mice were immunized in the flank by subcutaneous (s.c.) injection of 200 μg MOG_30-55_ peptide (MEVGWYRSPFSRVVHLYRNGK) (Peptide Synthesis Core Facilities of the Pompeu Fabra University, Barcelona, Spain) in incomplete Freund’s adjuvant supplemented with 8 mg/ml Mycobacterium tuberculosis H37Ra (Difco Laboratories). Pertussis toxin (500 ng; Calbiochem) was injected i.p. on the day of immunization and again 2 days later. Body weight and motor symptoms were recorded daily and scored from 0 to 8 as follows: 0, no detectable changes in muscle tone and motor behavior; 1, flaccid tail; 2, paralyzed tail; 3, impairment or loss of muscle tone in hindlimbs; 4, hindlimb hemiparalysis; 5, complete hindlimb paralysis; 6, complete hindlimb paralysis and loss of muscle tone in forelimbs; 7, tetraplegia; and 8, moribund.

### Cell culture

Primary astroglial cultures were prepared from homozygote CB_1_-WT by magnetic activated cell sorting (MACS). Briefly, the forebrains were carefully dissected, meninges removed and tissue enzymatically dissociated using Neural Tissue Dissociation Kit (P) (Miltenyi Biotec). Myelin was removed using a 35% Percoll PLUS (Merck) gradient and cells were sorted using Anti-ACSA-2^+^ MicroBead Kit (Miltenyi Biotec). Cells were seeded onto 14 mm diameter glass coverslips in 24-well plates at a density of 20.000 cells/well for immunocytochemistry and in 6-well plates at a density of 500.000/well for qPCR. Alternatively, astrocytes were seeded onto glass-bottom µ- dishes (Ibidi GmbH) at a density of 100.000 cells/plate for calcium imaging. Cells were maintained in serum-free defined medium containing 50% neurobasal, 50% DMEM, 100 U/ml penicillin, 100 μg/ml streptomycin, 1 mM sodium pyruvate, 292 μg/ml l-glutamine, 1×SATO and 5 μg/ml of *N*-acetyl cysteine. This medium was supplemented with the astrocyte-required survival factor HBEGF (10 ng/ml; Preprotech). The purity of the cultures was be routinely assessed by examining the characteristic cell morphologies under phase-contrast microscopy and confirmed by immunostaining with mouse anti-GFAP (1:200; Merck, MAB3402) and mouse anti-S100β (1:400; Sigma Aldrich, S2532). After 6-7 days in culture GFAP^+^ and S100β^+^ cells represented 95 ± 1% and 97 ± 2% of total cells, respectively (*n* = 3 cultures, 1-2 coverslips per culture, 15 microscopic fields per coverslip). Astrocyte activation to a neurotoxic phenotype was induced in cells at 4-5 DIV by incubation with TNFα (25 ng/ml; Cell Signaling Technology), IL-1α (3 ng/ml; Sigma), and C1q (400 ng/ml; MyBioSource) for 18-24 h (Liddelow et al., 2017) and corroborated by immunofluorescence and qPCR analysis (**Supplementary Fig.7**).

### Live cell imaging and data analysis

Astrocytes were loaded with Fluo-4 AM (1 mM; Molecular Probes) in culture media for 20 min at 37°C followed by 10 min wash in HBSS (Sigma, H-4891) supplemented with 20 mM HEPES, 10 mM glucose, 2 mM CaCl_2_, 1 mM MgCl_2_ and 4 mM NaHCO_3_ (pH 7.4) to allow de-esterification. Images were acquired through a 63X objective in a TCS SP8X STED CW confocal microscope (Leica) at an acquisition rate of 1 frame/10 s during 5 min. Following a baseline recording (1 min) cells were stimulated with WIN (1 µM), ATP (200 µM), glutamate (200 µM), (S)DHPG (100 µM), LY354740 (100 µM) or thapsigargin (1 µM). Recordings aimed at testing mitochondrial and/or endoplasmic reticulum responses were carried out in the absence of extracellular calcium. For data analysis, a homogeneous population of 10-15 cells was selected in the field of view and regions of interest (ROIs) defined within astrocyte somata. Background values were always subtracted and data are expressed as F/F_0_ in which F represents the fluorescence value for a given time point and F_0_ represents the mean of the resting fluorescence level.

### Surgery for AAV administration and fiber implantation

Mice (7-8 weeks of age) were anaesthetized with isoflurane and placed on a heating-pad to keep the body temperature at 37°C. Eye dehydration was prevented by topical application of ophthalmic gel and analgesia was achieved by s.c. injection of buprenorphine (Buprecare, 0.05 mg/kg). The skin above the skull was shaved with a razor and disinfected with modified ethanol 70% and betadine before an incision was made. Mice were placed into a stereotaxic apparatus (David Kopf Instruments) with mouse adaptor and lateral ear bars.

C57BL6N, IP_3_R_2_-KO and CB ^f/f^ mice were injected with AAV5-pZac2.1-gfaABC1d-cyto-GCaMP6f (Addgene), AAV-5/2-hGFAP-hM3D(G_q_)/hM4D(G_i_)_mCherry-WPRE-hGHp(A) (ETH Zurich) alone or in combination with and ssAAV-9/2-hGFAP-mCherry_iCre-WPRE-hGHp(A) (ETH Zurich) or ssAAV-9/2-hGFAP-mCherry-WPRE-hGHp(A) (ETH Zurich) for *ex vivo* imaging and electrophysiology. Virus v275-9 ssAAV- 9/2-GFAP-hHBbI/E-GCaMP6f-bGHp(A) was used for fiber photometry imaging of astrocytes in C57BL6N, CB_1_-KO and GFAP-CB_1_-KO mutant mice and control littermates. Virus titers were between 10^10^-10^12^ genomic copies per ml. Stereotaxic injections were targeted to the mouse somatosensory cortex according to the following coordinates (from bregma): anterior-posterior -1.5; medial-lateral ±2.5; dorsal-ventral - 1.5. Viral particles were injected at 400-500 nL alone or in combination at a maximum rate of 100 nL/min using a glass pipette attached to a Nanojet III (Drummond, Broomall, USA). Following virus delivery, the syringe was left in place for 10 min before being slowly withdrawn from the brain. For fiber photometry experiments, the optical fiber (400 μm diameter) was placed 250 µm above the injection site during the same surgical session. Mice were weighed daily and individuals that failed to return to their pre-surgery body weight were excluded from subsequent experiments. Wild-type, CB_1_- WT and CB_1_-KO mice and CB ^f/f^ mice were used for EAE induction 2 weeks after surgery. GFAP-CB_1_-KO and GFAP-CB_1_-WT mice were treated with tamoxifen 1 week after the surgery and were used for EAE experiments 3 weeks after the last tamoxifen injection.

### Cortical slice preparation for imaging and electrophysiology

Mice were euthanized by decapitation and brains were rapidly removed and placed in ice-cold artificial cerebrospinal fluid (aCSF). Coronal brain slices (350 µM-thick) containing the somatosensory cortex were prepared via a Leica VT1200 vibratome in a 4°C aCSF solution. Following cutting, slices were allowed to recover in aCSF containing (in mM): NaCl 124, KCl 2.69, KH_2_PO_4_ 1.25, MgSO_4_ 2, NaHCO_3_ 26, CaCl_2_ 2 and glucose 10, gassed with 95% O_2_/5% CO_2_ (pH = 7.3-7.4) at 31°C for 30 min followed by at least 30 min at 20-22°C before recording. Slices were then transferred to an immersion recording chamber and superfused at 2 ml/min with gassed aCSF and the temperature of the bath solution was kept at 34°C with a temperature controller TC- 324C (Warner Instruments Co.). In experiments aimed at testing the effects of pro-inflammatory mediators on astrocyte calcium dynamics slices were incubated with TNFα (25 ng/ml), IL-1α (3 ng/ml), and C1q (400 ng/ml) for 30 min immediately before recording.

### *Ex vivo* two-photon imaging

Two-photon microscopy imaging of layer V-VI astrocytes in the somatosensory cortex was performed using a Femto-2D microscope equipped with a tunable Ti:Sapphire MaiTai DeepSee laser (Spectra Physics) and GaAsP PMT detectors. The laser was tuned at 920 nm with a 20X water immersion lens (1.00 N.A.; Olympus) and a 490/60 nm filter. All calcium experiments were performed in the presence of TTX (1 µM). Videos were obtained at 512 × 512 resolution with a sampling interval of 1 s. A custom MATLAB program (Calsee: https://www.araquelab.com/code/) was used to quantify fluorescence levels in astrocytes. Calcium variations recorded at the soma and processes of the cells were estimated as changes of the fluorescence signal over baseline (ΔF/F_0_), and cells were considered to show a calcium event when the ΔF/F_0_ increase was at least two times the standard deviation of the baseline. For representative images and videos of astrocytic calcium activity an event-based analysis tool (Automatic Quantification and Analysis (AQuA): https://github.com/yu-lab-vt/AQuA) was used.

Basal astrocyte calcium activity was quantified from the astrocyte calcium event frequency, which was calculated from the number of events per min within 3 min of recording. Evoked astrocyte calcium activity was quantified from the calcium event probability, which was calculated from the number of calcium elevations grouped in 15 s bins recorded from 8-30 astrocytes per field of view. The time of occurrence was considered at the onset of the calcium event. For each astrocyte analyzed, values of 0 and 1 were assigned for bins showing either no response or a calcium event, respectively, and the calcium event probability was obtained by dividing the number of astrocytes showing an event at each time bin by the total number of monitored astrocytes (Navarrete & Araque, 2010). All the astrocytes that showed a calcium event during the experiment were used for the analysis. The calcium event probability was calculated in each slice, and for statistical analysis, the sample size corresponded to the number of slices. To examine the difference in calcium event probability in distinct conditions, the basal calcium event probability (mean of the 1 min before a stimulus) was averaged and compared to the average calcium event probability (15 s after a stimulus). For drug application, a micropipette was filled with WIN (300 µM), glutamate (200 µM), ATP (200 µM), (S)DHPG (100 µM), LY354740 (100 µM) or CNO (1 mM) solution and placed 100-150 µm away from the tissue and a pressure pulse at 1 bar was applied for 5 s. The effect of the CB_1_ antagonist AM251 (2 µM) was tested after 10 min bath perfusion in the same region and same astrocytes recorded in control conditions.

### Electrophysiology

Electrophysiological recordings from layer V pyramidal neurons were made using the whole-cell-patch-clamp technique. Cells were visualized using infrared-differential interference contrast optics (Olympus BX51WI microscope, Olympus Optical, Japan) and 40x water immersion lens and selected for recording based on their location, morphology, and firing pattern. When filled with an internal solution containing (in mM): 110 potassium gluconate, 40 HEPES, 4 NaCl, 4 ATP-Mg, and 0.3 GTP (pH = 7.3) patch electrodes exhibited a resistance of 3-10 MΩ. All recordings will be performed using MultiClamp 700B amplifier (Molecular Devices, San Jose, CA). Fast and slow whole-cell capacitances were neutralized, and series resistance was compensated (∼70%), and the membrane potential was held at -70 mV. Intrinsic electrophysiological properties were monitored at the beginning and the end of the experiments. Series and input resistances were monitored throughout the experiment using a -5 mV pulse. Recordings were considered stable when the series and input resistances, resting membrane, and stimulus artifact duration did not change > 20%. Signals were fed to a Pentium-based PC through a DigiData 1550 interface board. Signals were filtered at 1 kHz and acquired at a 10 kHz sampling rate using a DigiData 1550 data acquisition system and pCLAMP 10.3 software (Molecular Devices, San Jose, CA). Theta capillaries (2-5 µm tip) filled with ACSF were used for bipolar stimulation and placed in layer 2/3. Electrical pulses were supplied by a stimulus isolation unit (ISO-Flex, A.M.P.I, Jerusalem, Israel). Excitatory postsynaptic currents (EPSCs) were induced using brief current pulses (1 ms) delivered at 0.33 Hz and isolated using picrotoxin (50 µM) and CGP5462 (1 µM) to block GABA_A_ and GABA_B_ receptors, respectively. EPSC amplitude was determined as the peak current amplitude (2-20 ms after stimulus) minus the mean baseline current (10-30 ms before stimulus).

Input-output curves of EPSCs were obtained by increasing stimulus intensities from 0 to 100 μA in 10 μA steps. Paired-pulse facilitation was analyzed by applying paired pulses with 25, 50, 75, 100, 150, 200, 300 and 500 ms inter-pulse intervals. The paired-pulse ratio was calculated by dividing the amplitude of the second EPSC by the first (PPR = EPSC-2/EPSC-1). Synaptic fatigue was assessed by applying 19 consecutive stimuli in 25 ms intervals. AMPA and NMDA currents were obtained with an internal solution containing (in mM): cesium gluconate 117, HEPES 20, EGTA 0.4, NaCl 2.8, TEA-Cl 5, ATP 2, GTP 0.3. AMPA currents were obtained at a holding potential of −70 mV and NMDA currents at +30 mV. To ascertain the AMPA to NMDA receptor current ratio, we measured the NMDA component 50 ms after the stimulus.

For paired recordings, neurons were selected following the same vertical axis and distance of the somas of the layer V paired-recorded neurons varied from 70 to 270 µm. To induce endocannabinoid-mediated short-term synaptic plasticity (Baraibar et al., 2022) pyramidal neurons were depolarized from −70 mV to 0 mV for 5 s (ND). Synaptic parameters were determined from 40 stimuli before (basal) and following ND. Baseline mean EPSC amplitude was obtained by averaging mean values calculated within 2 min of baseline recordings and mean EPSC amplitudes were normalized to baseline. The ND was applied 2.5 s after the last basal delivered pulse, and no pulses were presented during the ND. The 0.33-Hz pulse protocol was restarted immediately after the ND step. To illustrate the time course of ND-induced effects, synaptic parameters were grouped in 30 s bins. Two consecutive responses to ND were averaged. For every synaptic recording, the presence of heteroneuronal depression or potentiation was assessed in individual synaptic recordings if the EPSC amplitude decreased or increased >2 times the standard deviation of the baseline EPSC amplitude during the first 45 s after the ND.

To induce spike-timing dependent plasticity (STDP) layer V pyramidal neurons were kept in voltage-clamp mode before and after the pairing protocol and EPSCs evoked by stimulation in layer II-III at 0.33-Hz. During the pairing protocol neurons were kept in current clamp mode and EPSPs were paired with one postsynaptic action potential at a time interval of Δt = −25 ms and pairings were repeated 60 times at 0.1 Hz (Min & Nevian, 2012). Experiments were discarded if the pyramidal neuron input resistance changed by >20% during the course of the experiment.

Miniature excitatory postsynaptic currents (mEPSCs) and slow inward currents (SICs) mediated by astrocytic glutamate release (Fellin et al., 2004; Perea & Araque, 2005; Pirttimaki et al., 2017) were isolated in the presence of tetrodotoxin (TTX, 1 µM), picrotoxin (50 µM), and CGP54626 (1 µM) to block voltage-gated Na^+^ channels, GABA_A,_ and GABA_B_ receptors, respectively. In all cases the effects of pharmacological agents were tested after 10 min bath perfusion and at < 40 min after entering whole-cell mode in the stimulating neuron.

### Fiber photometry imaging

Freely moving mice were imaged using 470 and 405 nm LEDs to excite GCaMP6s after 3 days of handling habituation. The emitted fluorescence is proportional to the calcium concentration for stimulation at 470 nm (Akerboom et al., 2013; Ohkura et al., 2012). The isosbestic 405 nm stimulation (UV light) was used in alternation with the blue light (470 nm) for analysis purposes as the fluorescence emitted after this stimulation is not depending on calcium (Lütcke et al., 2010). The GCaMP6s fluorescence from the astrocytes was collected with a sCMOS camera through an optic fiber divided in 2 sections: a short fiber implanted in the brain of the mouse and a long fiber (modified patchcord), both connected through a ferrule-ferrule (1.25 mm) connection. MATLAB program (Matlabworks) was used to synchronize each image recording made by the camera, and the Mito-GCaMP6s light excitation made by the LEDs (470 and 405 nm). The two wavelengths of 470 and 405 nm at a power of 0.1 mW were alternated at a frequency of 20 Hz each (40 Hz alternated light stimulations).

To calculate fluorescence due specifically to calcium fluctuations and to remove bleaching and movement artifacts, the isosbestic 405 nm signal was subtracted from the 470 nm calcium signal. Specifically, normalized fluorescence changes (Δ*F*/*F_0_*) were calculated by subtracting the mean fluorescence (2 min sliding window average) from the fluorescence recorded by the fiber at each time point and dividing this value by the mean fluorescence ((*F-F*_mean_)/*F*_mean_) using a customized Matlab software. Subsequently, the calcium independent isosbestic signal was subtracted to the raw signal emitted after the 470 nm excitation to eliminate unspecific fluorescence. The result will be the global Ca^2+^ signal (Δ*F*/*F* (%) = Δ*F*_Ca_ - Δ*F*_isos_), that was used as an estimate of tonic activity of the astrocytes. Ca^2+^ transients were detected on the filtered trace (high filter) using a threshold to identify them (2 median absolute deviation -MAD-of the entire trace). Duration and frequency were calculated on the detected transients. Amplitude was determined as the MAD of each studied period. The effects of vehicle and THC injection were analyzed in 2 equal time periods after injection (*period 1* and *period 2*).

The day of recording each mouse was placed in a rectangular chamber and its behavior recorded using a camera placed above the chamber. In experiments aimed at evaluating the effect of THC in naïve mice, baseline recordings were made for 12-20 min. Animals were subsequently injected i.p. with vehicle solution and recorded for further 30 min after injection (15 min *period 1* + 15 min *period 2*). Mice were returned to the home cage following vehicle recordings. At least 24 h after vehicle recordings mice were injected with THC solution (10 mg/Kg; i.p.) following a 12-20 min baseline recording period and imaged for the same time-windows. In experiments designed to address changes in astrocytic Ca^2+^ dynamics during autoimmune demyelination EAE was induced following the habituation period and recordings made for 12 min every 2 days starting 3 days before MOG administration. At the end of the experiment, mice were injected with vehicle and THC (10 mg/Kg; i.p.) administered in consecutive days (20-21 dpi) and recordings made as described above.

### Flow cytometry

Mice were decapitated under isoflurane anesthesia (IsoVet^®^, B Braun) and cells purified from the forebrain according to previously described procedures (Moreno-García et al., 2020)(Manterola et al. 2018). Briefly, tissue was dissected and placed in enzymatic solution (116 mM NaCl, 5.4 mM KCl, 26 mM NaHCO_3_, 1 mM NaH_2_PO_4_, 1.5 mM CaCl_2_, 1 mM MgSO_4_, 0.5 mM EDTA, 25 mM glucose, 1 mM L-cysteine) with papain (3 U/mL) and DNAse I (150 U/μL, Invitrogen) for digestion during 25 min at 37**°**C. Halfway through the incubation, the minced tissue was triturated 10 times using 5-ml serological pipettes. Following enzymatic digestion, cells were mechanically released by gentle passage through 23 G, 25 G and 27 G syringe needles. After homogenization, tissue clogs were removed by filtering the cell suspension through prewetted 40 μm cell strainers (Fisherbrand^TM^) to a 50 mL Falcon tube quenched by 5 mL of 20% heat inactivated fetal bovine serum in Hank´s Balanced Salt Solution (HBSS, Thermo Fisher Scientific). The cell strainers were thoroughly rinsed with 15 ml HBSS in and cell suspensions were centrifuged 200 x *g* for 5 minutes. To purify astrocytes from myelin debris, cells were resuspended in 25% isotonic Percoll PLUS (GE Healthcare Europe GmbH) in HBSS and centrifuged at 200 x *g* without brake for 20 min at room temperature. The myelin top layer was aspirated and cells washed with HBSS to remove any traces of Percoll PLUS by centrifuging at 200 x *g* for 5 min. The total dissociated single cells were resuspended in 500 μL sorting buffer (25 mM HEPES, 5 mM EDTA, 1% BSA, in HBSS) containing Normal Rat Serum (1:100; Invitrogen, 10710C) and TruStain FcX™ (anti-mouse CD16/32) antibody (1:100; BioLegend, 101320). Isolated cells were incubated with fluorochrome conjugated antibodies ACSA-2-PE (1:50; Miltenyi Biotec, REA969), A2B5-488 (1:100; R&D Systems, FAB1416G), CD11b-FITC (1:200; BioLegend, 101205), CD31-VioBright 515 (1:200; Miltenyi Biotec, REA784), CD45-FITC (1:200; Miltenyi Biotec, REA737), O1-488 (1:100; R&D Systems, FAB1327G) and LIVE/DEAD Fixable Green Dead Cell (1:1500; Thermo Fisher, L34969). Samples were run on a BD FACS Jazz (2B/4YG) flow cytometer (BD Bioscience) controlled using BD FACS™ Software (version 1.1.0.84) and results were analyzed using FlowJo software. All gatings were set based on appropriate isotype controls. Debris and cell clumps were initially gated out on the basis of FSC and SSC plots, allowing selection of only the population of interest. Further doublets were gated out using FSC/trigger pulse width plots. Astrocytes were sorted as ACSA-2^+^ cells following exclusion of contaminating microglia, hematopoietic cells, oligodendrocytes, oligodendrocyte progenitors and endothelial cells grouped in a dump channel. Typically, an average number of 50.000 astrocytes were collected for each sample within 15-20 min. The purity of astrocytes was assessed by flow cytometry analysis of sorted ACSA-2^+^ cells following fixation and immunostaining with a C-terminal antibody against the astrocytic marker GLAST (Rothstein et al., 1994) (1:200; kindly donated by Rothstein). We further confirmed that we had isolated a relatively pure population of astrocytes and microglia by qPCR analysis for the expression of several cell-type specific markers (Moreno-García et al., 2020).

### Quantitative RT-PCR

Anesthetized mice were transcardially perfused with cold phosphate buffer saline (PBS) for 30 s in order to remove blood cells from the brain. The somatosensory cortex was dissected from a single coronal slice trimmed between levels 1 and −1 bregma. Astrocytes purified by flow cytometry and cultured cells were collected in lysis buffer (Qiagen) containing 1% β-mercaptoethanol for optimal template preservation. Total RNA from brain samples, purified astrocytes and cultured cells was purified with on-column DNAse treatment using RNeasy Plus Micro and Mini kits (Qiagen) following manufacturer’s instructions. RNA was eluted with 14-35 μl of RNAse-free deionized water and stored at -80°C until analysis. Synthesis of cDNA, pre-amplification and amplification steps were performed at the Genome Analysis Platform of the UPV/EHU following quality control of RNA samples with an Agilent 2100 Bioanalyzer (Agilent Technologies). Pre-amplified cDNA samples were measured with no reverse transcriptase and no template controls in the BioMark HD Real-Time PCR System using 48.48 Dynamic Arrays of integrated fluidic circuits (Fluidigm Corporation). We used commercial primers from IDT Integrated DNA Technologies or Fluidigm Corporation (**Supplementary Table 1**). Data pre-processing and analysis were completed using Fluidigm Melting Curve Analysis Software 1.1.0 and Real-time PCR Analysis Software 2.1.1 (Fluidigm Corporation) to determine valid PCR reactions. *Gapdh*, *Hprt*, *Ppia* and *B2m* were included as candidate reference genes for normalization purposes. Data were corrected for differences in input RNA using the geometric mean of reference genes selected according to results from the normalization algorithms geNorm (https://genorm.cmgg.be/) and Normfinder (https://moma.dk/normfinder-software). Relative expression values were calculated with the 2^-ΔΔCt^ method.

### Fluorescence immunohistochemistry

Mice were anesthetized with chloral hydrate (400 mg/Kg) and transcardially perfused with 0.9% NaCl followed by 4% paraformaldehyde dissolved in 0.1 M phosphate buffer (PB; 25 mM NaH_2_PO_4_•H_2_O; 75 mM Na_2_HPO_4_; pH 7.4). After extraction, the brains were incubated for 3 hours in the same fixative and stored in PB at 4°C. Serial brain coronal sections between bregma -0.95 and -1.91 levels were cut at 40 µm and collected in PB at room temperature (RT) and stored in PB containing 0.02% sodium azide at 4°C until use.

For immunohistochemistry, floating sections were washed three times in Tris-HCl buffered saline (TBS; 100 mM Tris Base, 150 mM NaCl; pH 7.4) for 10 min and incubated in a blocking-permeabilization solution containing 5% normal goat serum and 0.2% Triton X-100 in TBS for 60 min at RT. Sections were subsequently incubated with the corresponding primary antibodies diluted in TBS supplemented with 1% NGS and 0.1% Triton X-100 for 24-72 h at 4°C. The following primary antibodies were used: chicken anti-GFAP (1:500, Abcam, Ab4674), mouse anti-S100b (1:200, Sigma, S2532), rabbit anti-C3d Complement (1:1000; Dako, A0063), rabbit anti-Iba1 (1:500; Wako, 019-19741), rat anti-CD68 (1:100; Bio-Rad, MCA1957GA), mouse anti-MBP (1:1000, Biolegend, 808402), mouse anti-Olig2 (1:500, Millipore, MABN50), rabbit anti-NG2 (1:500; Millipore, AB5320), mouse anti-APC/CC1 (1:500; Millipore, OP80), mouse anti-Synaptophysin (1:1000; BioLegend, 837101), rabbit anti-Homer-1 (1:500; Synaptic Systems, 160003) and rat anti-CD3 (1:50; Bio-Rad, MCA772). Tissue samples run in parallel without primary antibodies were always included as internal controls. Sections were washed three times in TBS for 10 min each before incubation for 1-2 h at RT with the following Alexa Fluor secondary antibodies made in goat (1:500; Invitrogen): anti-rabbit Alexa 488 (A11008), anti-rabbit Alexa 647 (A21244), anti-chicken Alexa 647 (A21449), anti-rat Alexa 488 (A11006), anti-mouse Alexa 488 (A11001), anti-mouse Alexa 594 (A11005), anti-mouse Alexa 647 (A21235), anti-mouse IgG2a Alexa 594 (A21135), anti-mouse IgG2b Alexa 488 (A21141). Hoechst 33258 (5 µg/mL; Sigma-Aldrich, 861405) was used together with secondary antibodies for chromatin staining. Finally, sections were washed three times for 10 min in TBS, mounted, dried and a coverslip was added on top with ProLong Gold antifade reagent (P36930, Invitrogen by ThermoFisher).

For immunofluorescence related with fiber photometry experiments, sections were washed three times for 5 min in PBS (0.1 M, pH 7.4) and incubated with rabbit anti-GFP (1:500; Invitrogen, A11122) and chicken anti-GFAP (1:500; Abcam, ab4674) overnight at 4°C in a blocking solution containing 10% donkey serum and 0.3% Triton X-100 in PBS. The sections were then washed in PBS for 30 min at RT and primary antibodies were detected by incubation with donkey anti-rabbit Alexa 647 (1:500; Invitrogen, A31573) and donkey anti-chicken Rhodamine Red (1:500; Jackson ImmunoResearch, 2340371) for 2 h at RT. Then, sections were washed for 15 min in PBS, mounted, dried and mounted on microscope slides in Fluoromount G.

Optical images from tissue sections processed in parallel were acquired in the same session using a 20X objective lens and a 40X or a 63X oil immersion lens on a Zeiss Axioplan 2 pseudoconfocal microscope and Leica TCS STED CW SP8 super-resolution microscope. Image acquisition was carried out using fluorescence intensity settings at which the control sections without primary antibody gave no signal. Immunolabeling was examined bilaterally in 4 pictures per tissue section and analysed using Fiji Image J. Immunopositive cells were quantified by cell counting and data expressed as mean cell number per square millimeter (mm^2^) of tissue area. Threshold analysis of GFAP immunostained area was carried out in 16-bit grey scale transformed pictures and the values were referred to the specific layer area in each optical section. Custom scripts were used for semiautomated image processing and analysis of co-localization of GFAP+C3d and Iba1+CD68 as well as for synapse counting in Fiji Image J (Schindelin et al. 2012).

### Statistical analyses

Summary results are presented as mean of independent data points ± SEM that represent the number of animals, slices, neurons or cultures tested. Datasets were initially tested for normal distribution and statistical analysis of the differences between groups were determined by two-tailed unpaired or Student paired *t* test, Mann-Whitney test, one-way ANOVA or two-way ANOVA followed by Šídák’s test for multiple comparisons. Differences were considered to be significantly different when *P* < 0.05. Correlation analysis was performed by Pearson’s or Spearmańs test.

## Results

### Emergence of aberrant astrocyte calcium responses during EAE

To study the calcium activity of astrocytes during cortical MS pathology we used GCaMP6-based fiber photometry in freely behaving mice. As experimental model of MS we induced chronic EAE by immunization with myelin oligodendrocyte glycoprotein (MOG_35-55_) in mice injected GFAP-GCaMP6f viral particles (**Fig. 1a**). We monitored astrocytes in the somatosensory cortex during disease time-course starting 3 days before MOG_35-55_ administration. The first clinical manifestations occurred at 10 dpi with acute disease peaking at 18 dpi and attenuation of neurological severity taking place at later time points (**Fig. 1b**). Histological analysis of *postmortem* tissue at the experimental end-point showed specific expression of the calcium sensor in astrocytes from control and EAE mice (**Supplementary Fig. 1**).

**Figure 1.**
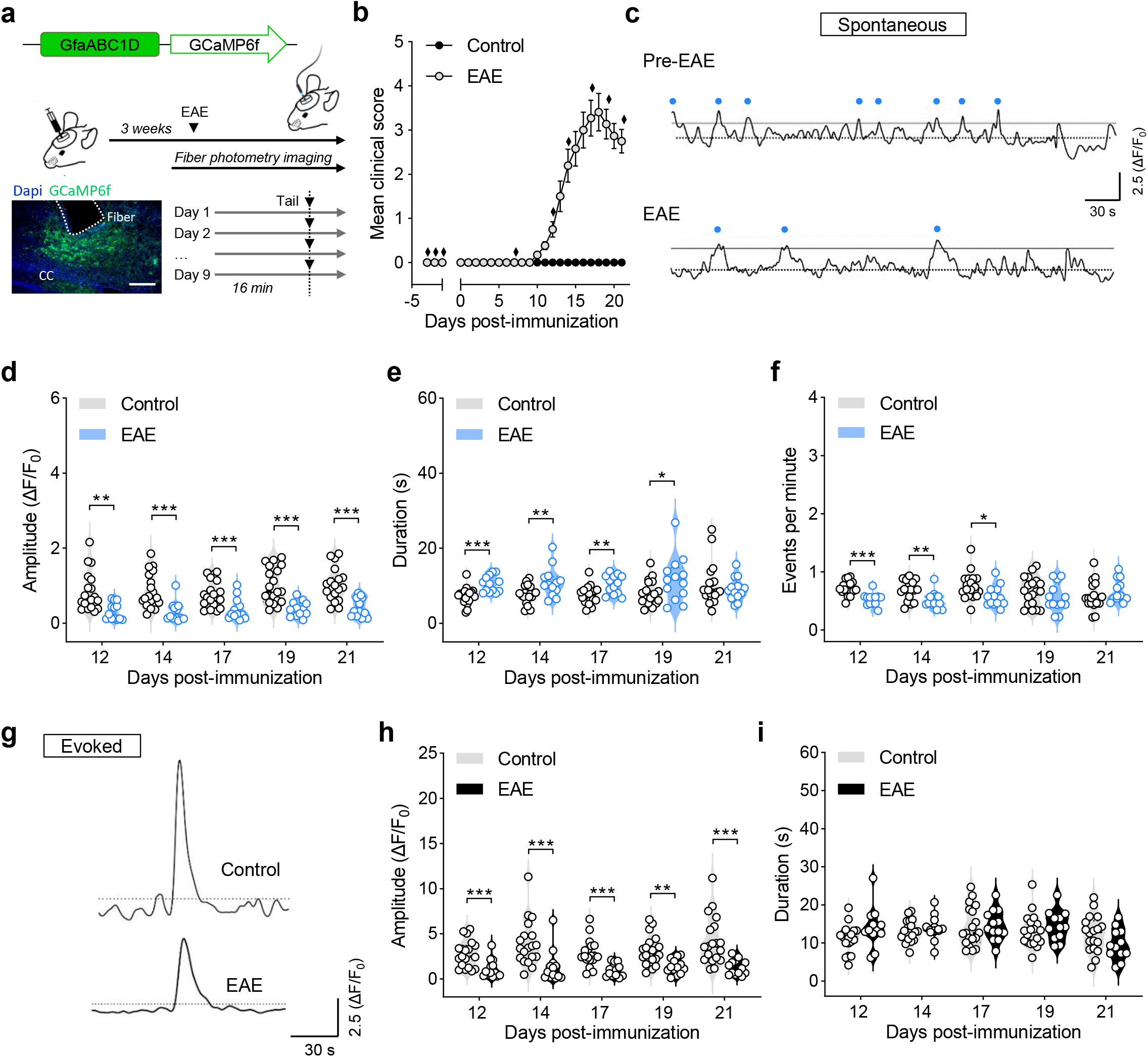
Emergence of aberrant astrocyte calcium activity in the somatosensory cortex of EAE mice. (**a**) Experimental fiber photometry approach for *in vivo* recording of astrocytic calcium activity during EAE time-course. Fiber photometry imaging was performed in control and EAE mice on 9 independent recording sessions as depicted in **b** (diamonds). Below: representative image of GCaMP6s expression in the somatosensory cortex. DAPI is marked in blue and GCaMP6s in green. Scale bar: 200 µm. (**b**) Neurological score of EAE mice included in the study (19 mice; 3 independent EAE experiments). Data are mean ± SEM. (**c**) Representative traces of spontaneous activity in cortical astrocytes during the pre-immunization phase (-1 dpi) (top) and during EAE disease (14 dpi) (bottom). Blue dots correspond to transients detected above the threshold (median+2*MAD). (**d**-**f**) Analysis of spontaneous astrocyte calcium oscillations in the somatosensory cortex of EAE mice at different time-points of disease progression as compared to naïve animals (19 mice). \**P* < 0.05; \*\**P* < 0.01; \*\*\**P* < 0.001; two-tailed unpaired *t* test or Mann-Whitney test. (**g**) Representative traces showing calcium responses of cortical astrocytes evoked by tail-holding in control (top) and EAE (17 dpi) (bottom) mice. (**h-i**) Amplitude and duration of evoked calcium transients evoked in control and immunized mice. \**P* < 0.05; \*\**P* < 0.01; \*\*\**P* < 0.001; two-tailed unpaired *t* test or Mann-Whitney test.

Cytosolic calcium in cortical astrocytes can occur both spontaneously as well as in response to exogenous stimuli such sensory stimulation or footshock (Qin et al., 2020; Stobart et al., 2018). To investigate the impact of autoimmune inflammation on the calcium activity of cortical astrocytes we first addressed possible changes in spontaneously occurring calcium fluctuations *in vivo*, which reflect the integration of activity-dependent and independent cellular signals (Wang et al., 2006; Zur Nieden & Deitmer, 2006). The onset of EAE symptomatology was associated to a progressive decline in the amplitude of spontaneous calcium transients in cortical astrocytes that reached a plateau at 14 dpi (**Fig. 1c, d**). Further analysis showed that spontaneous astrocyte calcium activity displays events of increased duration and reduced frequency in EAE mice as compared to control animals recorded in parallel (**Fig. 1c-f**). We next investigated the possible relationship between deregulated calcium responses in cortical astrocytes and motor deficits, which reflect spinal cord axon loss during EAE progression (Wujek et al., 2002). Noteworthy, reductions in the amplitude of spontaneous calcium transients in freely-behaving mice correlated to EAE neurological disability thus pointing to an association with spinal cord pathology (**Supplementary Fig. 2a**, top panel). However, changes in the duration and frequency of astroglial calcium signals did not correlate to the severity of motor deficits at any of the time-points tested (**Supplementary Fig. 2a**, middle and bottom panels). These observations suggest that deregulation of spontaneous calcium oscillations in cortical astrocytes involves deficits in the transmission of motor signals associated to the emergence of spinal cord inflammatory lesions during EAE as well as local, autocrine and/or paracrine cellular interactions within the somatosensory cortex. To further investigate this issue we addressed changes in astrocytic calcium activity evoked by sensory stimulation of the tail in EAE mice. Under our experimental settings, suspension by the tail induced reliable calcium responses in cortical astrocytes from naive mice that remained stable in size during the chronic recording period (**Fig. 1g**). EAE induction was associated to a significantly decreased amplitude of astrocytic calcium responses evoked by tail-holding that paralleled symptom onset (**Fig. 1g**, **h**). By contrast, the duration of evoked astrocyte calcium signals remained unchanged during the time course of EAE. Reminiscent of our observations concerning spontaneous calcium responses, we observed an inverse correlation between the amplitude of astrocyte calcium responses evoked by tail-holding and neurological disability score values during EAE progression that reached statistical significance at 17 and 19 dpi (**Supplementary Fig. 2b**). Overall, these data suggest that deficits in the amplitude of spontaneous and activity-dependent astrocyte calcium oscillations in the motor cortex may reflect impairments in the conduction of signals that result from degeneration of spinal cord axons during EAE.

We next addressed the involvement of local mechanisms in the emergence of dysregulated astrocyte calcium activity during EAE using *ex vivo* two-photon microscopy of GCaMP6f in cortical slices. Experiments were performed in the presence of TTX to record activity-independent astrocyte oscillations and thus exclude potential confounding effects from altered cortical excitatory-inhibitory balance associated to EAE pathology (Potter et al., 2016). Under these experimental conditions, astrocytes within layer V-VI of the somatosensory cortex exhibited sparse spontaneous calcium activity in control conditions (**Fig. 2a**, **b**). We observed a 2-fold increase in the calcium event probability of cortical astrocytes at acute EAE disease (**Fig. 2a-c**; **Supplementary videos S1** and **S2**). Immunized mice displayed astrocyte calcium events of larger amplitude (**Fig. 2d**) and drastically enhanced duration (**Fig. 2b**; **Supplementary videos S3** and **S4**). Collectively, these results suggest that astrocytes of the somatosensory cortex display spontaneous calcium hyperactivity during EAE despite exhibiting impaired responses to sensorimotor stimuli *in vivo*.

**Figure 2.**
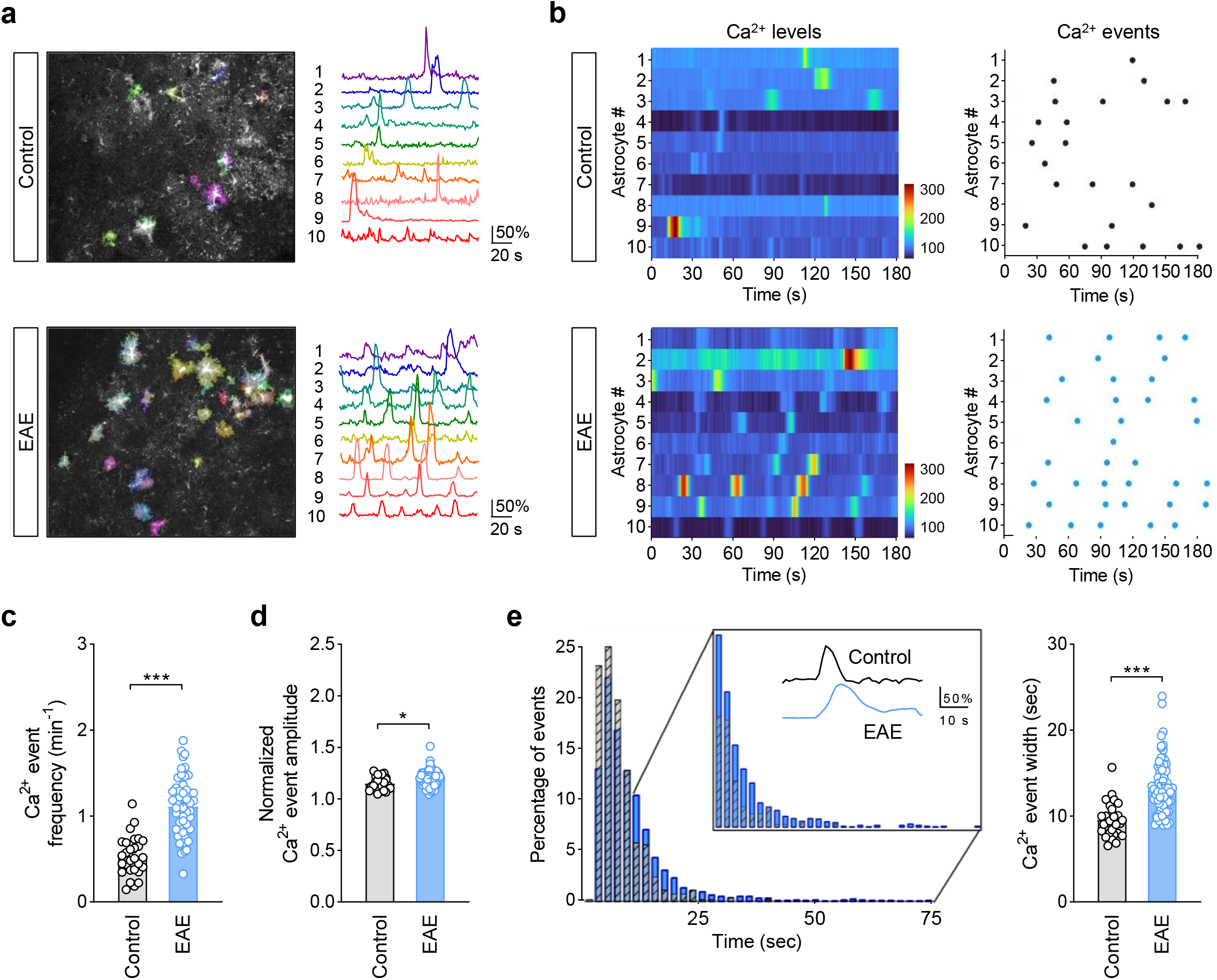
Spontaneous hyperactivity of cortical astrocytes during EAE. (**a**) Representative images and individual traces of calcium events accumulated over a 3 minute period in cortical astrocytes from control (top) and EAE (bottom) mice. (**b**) Heat maps and raster plots depicting calcium levels and spontaneous events along time in control and EAE astrocytes. (**c**) Frequency and amplitude of calcium events in cortical astrocytes from control (28 slices, 5 mice) and EAE mice (53 slices, 8 mice). (**d**) Left: Representative histogram showing the percentage of astrocyte calcium events and their duration in control and immunized mice. Box: percentages of events with longer duration in and representative traces from control (gray) and EAE (blue) astrocytes. \**P* < 0.05, \*\**P* < 0.01; \*\*\**P* < 0.001; two-tailed Student’s unpaired *t* test or Mann-Whitney test. Error bars express SEM.

### EAE impairs CB_1_ receptor-mediated astrocyte calcium oscillations

A prominent mechanism by which cortical astrocytes communicate with their environment is endocannabinoid signaling, which evokes intracellular calcium elevations through astrocyte CB_1_ receptors *ex vivo* and *in vivo* (Baraibar et al., 2022; Min & Nevian, 2012; Serrat et al., 2021). Changes in the levels of endocannabinoid ligands and receptor proteins have been pinpointed in human MS (Eljaschewitsch et al., 2006) and animal models of the disease (Centonze et al., 2007; Moreno-García et al., 2020) prompting us to explore potential alterations in astrocyte calcium responses evoked by CB_1_ receptors during EAE. To address this issue, we first performed fiber photometry recordings in naïve mice carrying GCaMP6f in cortical astrocytes treated with the plant-derived cannabinoid compound D^9^-tetrahydrocannabinol (THC, 10 mg/Kg) (**Supplementary Fig. 3a**). Time course image analysis revealed that THC treatment increased the amplitude and duration of astroglial calcium oscillations in the somatosensory cortex of naïve CB_1_-WT mice, but not in CB_1_-KO mice (**Supplementary Fig. 3b-d**). These results are consistent with our recent observation that CB_1_ receptor activation increases mitochondrial calcium in cortical astrocytes in freely-moving mice (Serrat et al., 2021). We next used conditional mutant mice lacking CB_1_Rs in GFAP-positive cells (GFAP-CB_1_-KO mice) (Han et al. 2012) to decipher whether the increase of astrocyte cytosolic calcium activity induced by THC involves the activation of CB_1_ receptors present in astroglial cells. Animals received tamoxifen injections 1 week after the surgery and were used for *in vivo* imaging experiments after a 3 weeks washout period (see methods for further details). Systemic THC did not modulate the dynamics of astroglial calcium events in the somatosensory cortex of GFAP-CB_1_-KO mice while reliably increased the amplitude of calcium signals in control littermates recorded in parallel (GFAP-CB_1_-WT) (**Supplementary Fig. 3f-g**). Thus, THC increases astrocyte calcium activity in the somatosensory cortex of freely-behaving mice through the activation of CB_1_ receptor populations present in astroglial cells.

Next, we explored whether the observed changes in the activity patterns of astrocytic calcium responses during EAE involved deficient regulation of cytosolic calcium levels by CB_1_ receptors. Mice subjected to EAE were sequentially challenged with vehicle and THC at 20-21 dpi in 2 consecutive sessions separated 24 hours (**Fig. 3a**). Mirroring results in naïve CB_1_-WT mice (**Supplementary Fig. 3b-d**), systemically administered THC effectively increased the amplitude and duration of astroglial calcium transients during the second half of the recordings in control non-immunized mice chronically monitored over several weeks (**Fig. 3b-d**). Noteworthy, injection of THC did not modulate the occurrence of astroglial calcium oscillations in the somatosensory cortex of EAE mice during the recording session suggesting deficits in astrocyte CB_1_ receptor-mediated calcium responses associated to disease progression. To further corroborate this hypothesis, we monitored calcium responses to the cannabinoid agonist WIN55,212-2 (WIN, 300 μM) in cortical astrocytes of EAE mice using *ex vivo* two-photon microscopy in acute slices. While local application of WIN increased spontaneous astrocyte calcium activity, quantified from the calcium event probability in naïve mice, WIN-evoked responses were significantly reduced in EAE mice (**Fig. 4b-d**; **Supplementary videos S5** and **S6**). Thus, EAE is associated to deficits in the ability CB_1_ receptors to modulate cytosolic calcium oscillations in cortical astrocytes.

**Figure 3.**
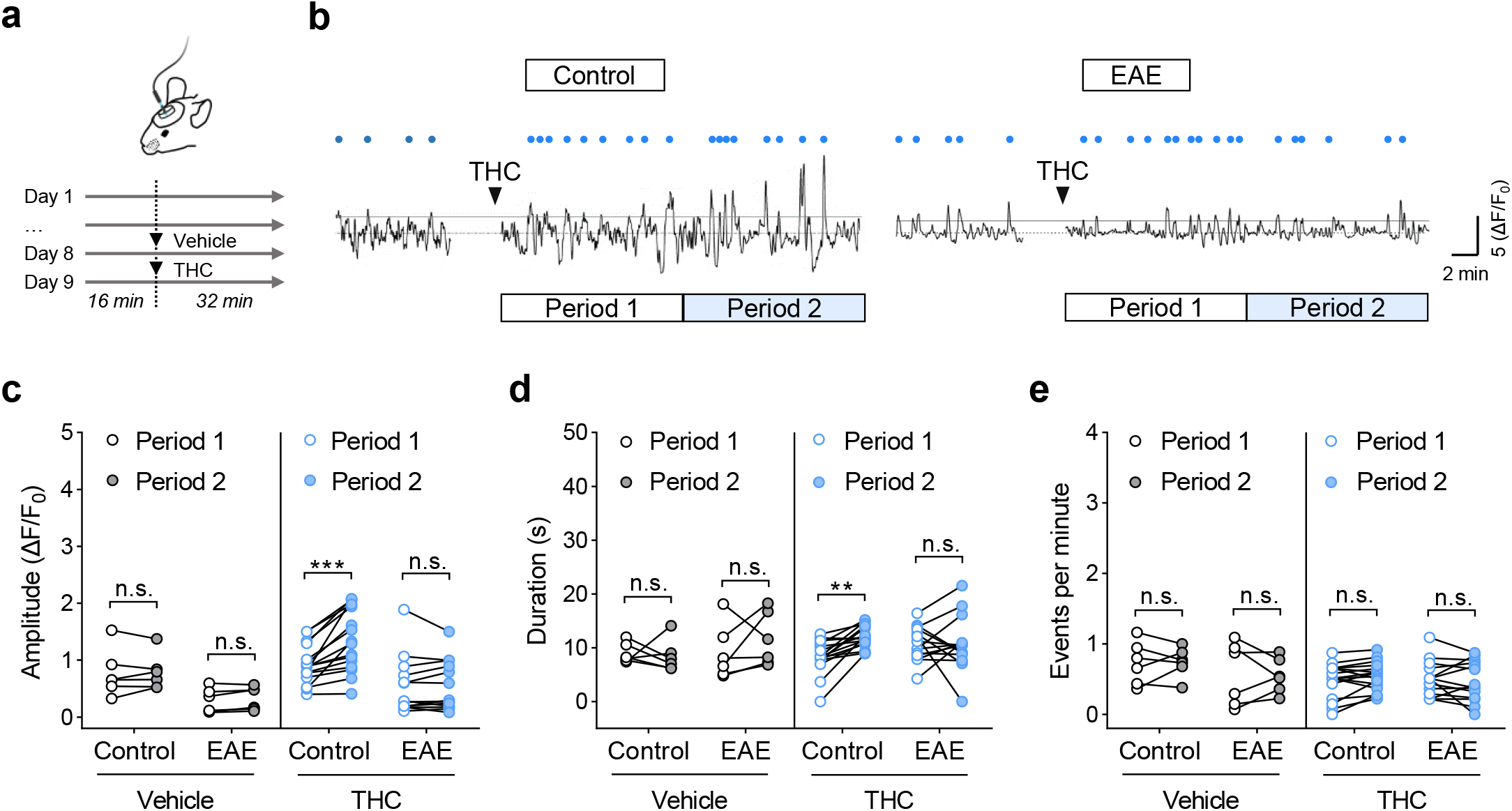
EAE impairs CB_1_ receptor-mediated astrocyte calcium responses. (**a**) Fiber photometry approach for *in vivo* recording of astrocytic calcium responses to THC in EAE mice. (**b**) Representative recordings from the mouse somatosensory cortex showing the effect of THC (10 mg/Kg; i.p.) on astrocytic calcium activity in control (left) and EAE (right) mice. Blue dots correspond to detected transients above the threshold (median+2*MAD) in mice injected with vehicle and THC. White and blue rectangles show the time window of analyzed period 1 and period 2, respectively. The first minute before and after injection was removed to exclude mouse-handling/injection effects. (**c**- **e**) Quantitative analysis of astrocyte calcium responses to vehicle and THC in control (6 mice) and EAE mice (17 mice). \*\**P* < 0.01; \*\*\**P* < 0.001; two-way ANOVA followed by Šídák’s multiple comparisons test. n.s., not statistically significant.

**Figure 4.**
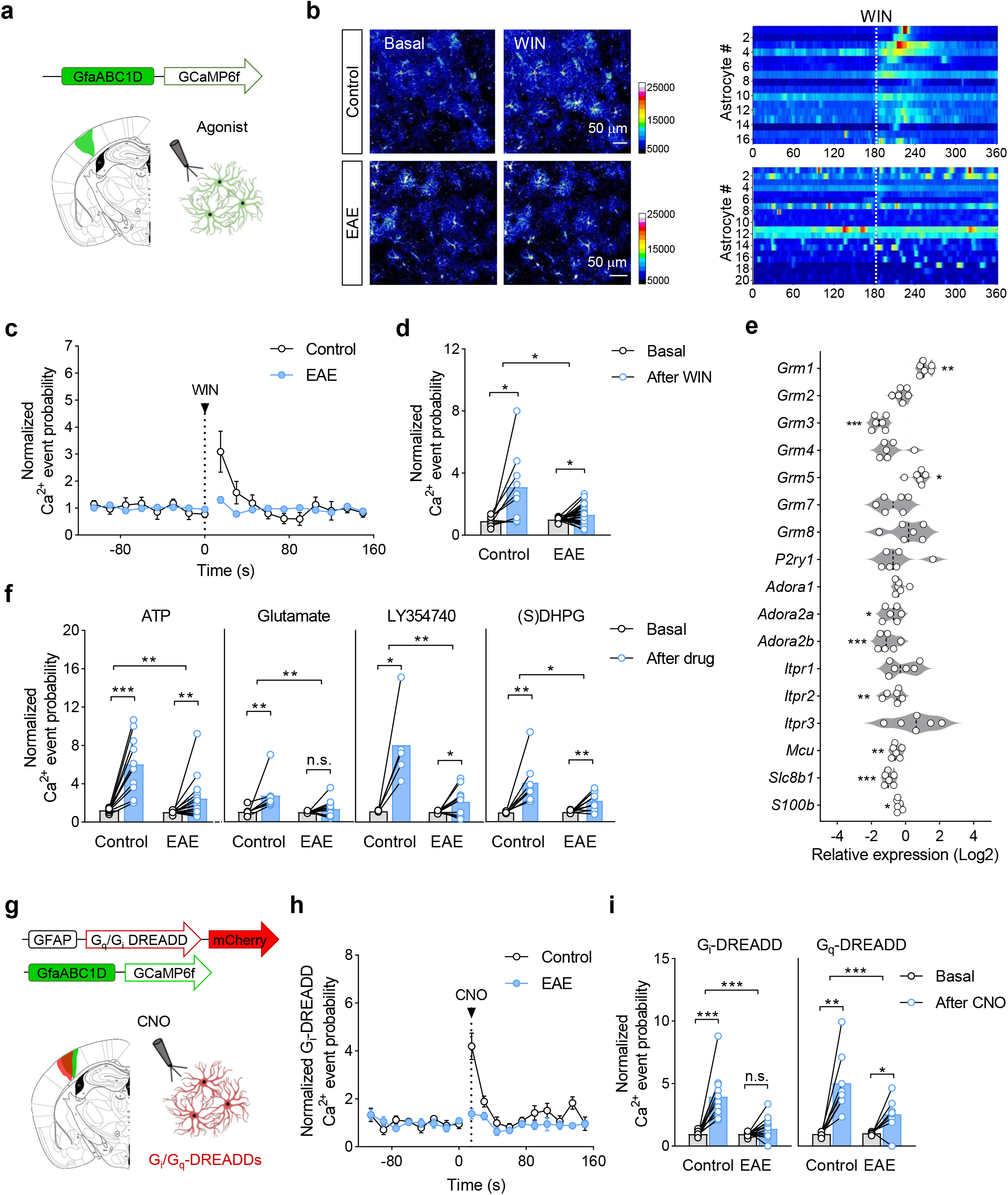
Cell autonomous deregulation of astrocyte calcium signaling pathways during EAE. (**a**) *Ex vivo* analysis of neurotransmitter receptor-mediated astrocyte calcium responses during EAE. Agonists were applied locally to the somatosensory cortex in acute slices from mice injected with the AAV vector encoding for GCaMP6f under the Gfa-ABC1D promoter. (**b**) Left: pseudocolor images obtained from cortical astrocytes of control (top) and EAE (bottom) mice before and after local application of the CB_1_ receptor agonist WIN55,212-2 (WIN; 300 µM). Right: heat map depicting calcium levels along time following application of the cannabinoid agonist in control and EAE astrocytes. (**c**) Normalized astrocyte calcium event probability over time showing the effect of WIN in control and immunized mice. (**d**) Bar graph depicting calcium event probability normalized to basal activity before and after WIN application in control (9 slices, 5 mice) and EAE astrocytes (25 slices, 7 mice). The dotted lines in **b** and **c** indicate 5 sec of WIN application. \*\**P* < 0.05; two-tailed paired Student’s *t* test (before and after) and unpaired *t* test or Mann-Whitney test (between groups). (**e**) Expression levels of calcium handling genes in astrocytes purified at acute EAE disease relative to those in healthy controls. \**P* < 0.05, \*\**P* < 0.01; \*\*\**P* < 0.001; two-tailed unpaired Student’s *t* test or Mann-Whitney test. (**f**) Calcium event probability normalized to basal activity before and after application of ATP (200 µM; 11-15 slices, 5-7 mice), glutamate (200 µM; 8-13 slices, 4-6 mice), LY354740 (100 µM; 5-14 slices, 2-2 mice), (S)DHPG (100 µM; 8-10 slices, 2-2 mice) in control and EAE astrocytes. \**P* < 0.05, \*\**P* < 0.01; \*\*\**P* < 0.001; two-tailed paired Student’s *t* test (before and after) and unpaired t test or Mann-Whitney test (between groups). (**g**) *Ex vivo* analysis of G_i_ and G_q_ mediated signaling on astrocyte calcium responses during EAE. Clozapine-N-oxide (CNO, 1 mM) was applied locally in acute cortical slices from mice injected with AAV vectors encoding for GCaMP6f and for G_i_ or G_q_ DREADDs under the astrocyte promoters GFAP or GfaABC1D. (**h**) Calcium event probability normalized to basal activity over time before and after CNO application in control and EAE mice injected with AAV8- GFAP-G_i_-DREADD-mCherry. The dotted line indicates 5 sec of CNO application. (**i**) Normalized calcium event before and after CNO application in control and EAE astrocytes expressing G_i_-DREADDs (11-15 slices; 2-4 mice) or G_q_-DREADDs (8-9 slices, 3-3 mice). \*\**P* < 0.01; \*\*\**P* < 0.001; two-tailed Student’s paired *t* test (before and after) and unpaired *t* test or Mann-Whitney test (between groups). n.s., not statistically significant. Error bars express SEM.

### Cell autonomous deregulation of astrocyte calcium dynamics involves deficits in G_i_ and G_q_ signaling pathways

Aberrant calcium responses of cortical astrocytes during EAE may be cell autonomous and/or reflect disease-associated changes in the dynamics of extracellular signals coupled to the modulation of calcium oscillations in these cells (i.e. endocannabinoids, glutamate or ATP/adenosine). Of note, astrocytes activated during EAE display reduced gene expression levels of CB_1_ receptors (Moreno-García et al., 2020), which provides a plausible cell autonomous mechanism underlying the observed deficits in calcium responses to cannabinoid agonists *ex vivo* and *in vivo*. Thus, we next asked whether reorganization of astrocyte calcium signaling during EAE involves abnormal expression of additional membrane receptors and calcium handling molecular cascades using RT-qPCR. Astrocytes purified from mice at acute EAE disease showed deregulated transcript levels of several calcium handling signaling/homeostatic toolkits (**Fig. 4e**). Among the genes whose expression were altered, we found complex changes in several receptors mediating glutamate signaling to astrocytes, such as metabotropic glutamate receptors (mGluRs) type 1 (*Grm1*), 3 (*Grm3*) and 5 (*Grm5*) (Haustein et al., 2014; Sun et al., 2013) as well as decreased levels of several adenosine receptors (*Adora2a*, *Adora2b*). Our RT-qPCR analysis also showed reductions in the expression levels of molecules chiefly involved the regulation of astrocytic calcium responses by the endoplasmic reticulum and the mitochondria, namely the inositol 1,4,5-trisphosphate (IP_3_) receptor type 2 (*Itpr2*) and the mitochondrial calcium uniporter (*Mcu*). Altogether, these results suggest that astrocytes at acute EAE disease display widespread deficits in calcium signaling pathways that are essential for their physiological functions.

To decipher whether deficits in astrocyte CB_1_ receptor-mediated calcium responses extend to signaling by additional neurotransmitter receptors we conducted two-photon imaging experiments in acute cortical slices from EAE mice. Calcium signals evoked by locally applied ATP and glutamate (200 μM) were significantly reduced at acute EAE disease thus mirroring impaired responses to the cannabinoid agonist WIN (**Fig. 4f**). We also found significantly reduced calcium responses to the mGluR_2/3_ agonist LY354740 (100 μM; **Fig. 4f**) which are consistent with the deficits in astrocyte *Grm3* expression during EAE determined by qPCR (**Fig. 4e**). Noteworthy, calcium oscillations evoked by the mGluR_1/5_ agonist (S)DHPG (300 μM) in cortical astrocytes was also slightly decreased (**Fig. 4f**) despite the augmented gene expression levels of both receptor proteins in cells purified from EAE mice (**Fig. 4e**). These combined results support the possibility that reorganization of astrocyte calcium signaling in the EAE cortex involves differential deficits in the intracellular pathways operated by G_i_ and G_q_ proteins in addition to deregulated expression of G protein-coupled receptors. To corroborate this hypothesis, we next investigated the functional consequences of selective activation of G_i_ and G_q_ signaling on astrocytic calcium activity using designer receptors exclusively activated by designed drugs (DREADDs). Astrocytes in the somatosensory cortex were specifically targeted with AAV8-GFAP-G_q_-DREADD-mCherry or AAV8-GFAP-G_i_-DREADD-mCherry in combination with AAV5-gfaABC1d-GCaMP6f and challenged with clozapine-N-oxide (CNO; 1 mM) during acute EAE (**Fig. 4g**). We found that application of CNO transiently elevated calcium levels in the soma and processes of cortical astrocytes expressing G_i_ or G_q_ DREADDs as quantitatively shown by the increase of the calcium event probability in naïve and EAE mice (**Fig. 4h**, **i**). These results confirm and extend previous observations on the ability of both signaling pathways to activate astroglial cells in term of calcium dynamics (Baraibar et al., 2022; Durkee et al., 2019). EAE induction completely abolished calcium oscillations induced by CNO in cells expressing G_i_ DREADDs (**Fig. 4h**, **i**). Responses to CNO were also significantly reduced, albeit to a lower extent, astrocytes from EAE mice expressing G_q_ DREADDs as compared to non-immunized animals (**Fig. 4i**). The G_i_ to G_q_ calcium signaling ratio calculated from the acute increase in calcium event probability following DREADDs activation was decreased by > 2-fold during EAE (control G_i_/G_q_: 74%; EAE G_i_/G_q_: 28%). These results provide evidence that cortical astrocytes exhibit disbalanced calcium oscillations mediated by G_i_ and G_q_ protein signaling pathways that may underlie, at least in part, abnormal calcium dynamics during EAE.

### Cortical pathology involves demyelination, inflammation and astrocyte reactivity

Cortical pathology in MS is characterized by inflammation dominated by innate immune cells, demyelination and a variable extent of synapse loss and neuronal death (Lagumersindez-Denis et al., 2017; Mahad et al., 2015). Previous studies have highlighted that the emergence of synaptic abnormalities in EAE mouse cortex precedes the onset of local inflammatory responses (Yang et al., 2013). Thus, to gain further insight into the cellular and molecular mechanisms that encompass aberrant astrocyte calcium signaling during EAE we next examined cortical neuropathology and synaptic function in the acute phase of the disease. Immunofluorescence staining for myelin basic protein (MBP) revealed that EAE mice display significant demyelination of the somatosensory cortex at 17 dpi (**Supplementary Fig. 4a-c**). We then immunostained for Olig2 to mark the oligodendrocyte lineage, in combination with CC1 or NG2 to identify myelinating oligodendrocytes or oligodendrocyte precursor cells, respectively. We observed a decline in the presence of Olig2^+^ oligodendroglial cells in cortical layers V-VI (411.1 ± 26.9 cells/mm^2^ EAE; 528.8 ± 23.4 cells/mm^2^ control; *P* = 0.0024; Student´s t test) that was associated to reduced numbers of mature oligodendrocytes and oligodendrocyte precursors (**Supplementary Fig. 4d-e**). Notably, there was a lower percentage of CC1^+^/Olig2^+^ mature oligodendrocytes and a higher percentage of NG2^+^/Olig2 oligodendrocyte precursors in the demyelinating EAE cortex (**Supplementary Fig. 4f**). Thus, astrocyte calcium signaling abnormalities encompass the loss of mature oligodendroglial cells likely underlying cortical demyelination during EAE.

Diminished neuronal activity and synapse loss preferentially affecting excitatory inputs are induced in rodent models of neuroinflammation that mimic cortical MS lesions (Jafari et al., 2021; Lagumersindez-Denis et al., 2017). Hence, we next sought to determine whether astrocyte dysfunction and cortical demyelination are associated to synaptic pathology at acute EAE disease. The density of excitatory synapses identified by immunostaining of the presynaptic synaptophysin and postsynaptic scaffold protein Homer-1 remained largely unchanged in the EAE cortex (**Supplementary Fig. 5a, b**). Consistently, we did not find differences between naïve and EAE mice regarding cortical mRNA expression of the neuron/synaptic markers such synapsin 1 (*Syn1*), syntaxin 1B (*Sytx1b*) and synaptosome associated protein 25 (*Snap25*) (**Fig. 5b**). Thus, cortical pathology at acute EAE disease does not appear to involve excitatory synapse loss. To further analyze the extent of synaptic pathology in the EAE mouse cortex we next evaluated the features of cortical excitatory synaptic transmission using *ex vivo* electrophysiology. The synaptic input-output relationship, the paired-pulse ratios and the AMPA/NMDA receptor relationship of EPSCs evoked in layer V cortical pyramidal neurons from mice at acute EAE did not differ from that in naïve animals suggesting intact excitatory transmission (**Supplementary Fig. 6a-d**). The frequency and the amplitude of miniature EPSCs (mEPSCs) was also nondiscriminable between control and EAE mice further indicating unaltered excitatory synaptic input (**Supplementary Fig. 6e**). Of note, mEPSCs in cortical neurons displayed increased duration associated to an augmented half-width and a slower rise and decay-phases (**Supplementary Fig. 6f**). These results are reminiscent of previous observations in the striatum and the cerebellum of EAE mice (Centonze et al., 2009; Mandolesi et al., 2013) and show that cortical astrocyte malfunction during EAE is encompassed by subtle changes in excitatory synaptic transmission.

**Figure 5.**
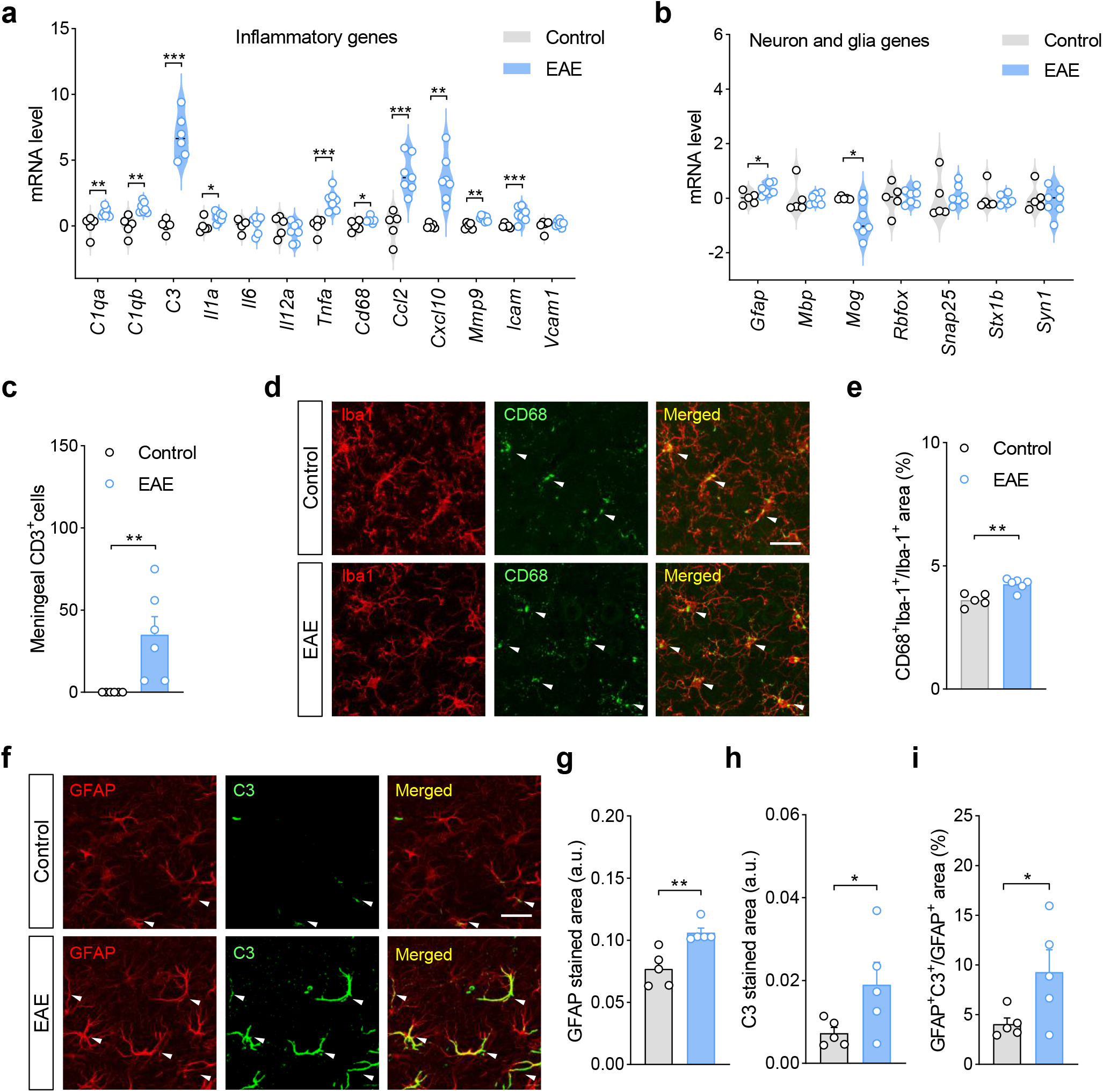
Local inflammation in the cortex of EAE mice. (**a**-**b**) Gene expression of pro-inflammatory mediators (**a**) and neuronal and glial cell markers (**b**) in the somatosensory cortex of EAE mice at acute disease (16 dpi; 7 mice) relative to those in control animals (5 mice). \**P* < 0.05, \*\**P* < 0.01; \*\*\**P* < 0.001; two-tailed paired Student’s *t* test or Mann Whitney test. (**c**) Quantitative analysis of meningeal CD3^+^ immunopositive T cells in coronal sections of EAE (14 dpi; 6 mice) and control conditions (5 mice). \*\**P* < 0.01; Mann Whitney test. (**d**) Representative confocal images showing the distribution of CD68 immunopositive puncta and Iba1-positive microglia in layer VI of the somatosensory cortex from control and EAE mice (17 dpi). Scale bar = 25 µm. (**e**) Quantification of CD68 levels within Iba1-positive profiles indicate microglia activation the EAE cortex (5-6 mice). \*\**P* < 0.01; two-tailed unpaired Student *t* test. (**f**) Representative confocal micrographs depict immunolabeling of the astrocyte reactivity marker GFAP and complement component 3 (C3) in layer VI from control and EAE conditions. Scale bar = 25 µm. (**g**-**i**) Quantitative analysis of GFAP (**g**) and C3 (**h**) levels and C3 expression in astroglial profiles (**i**) shows augmented astrocyte reactivity in the EAE cortex. a.u., arbitrary units. \**P* < 0.05, \*\**P* < 0.01; two-tailed unpaired Student *t* test. Error bars express SEM.

Because alterations of spontaneous glutamate transmission and synaptic plasticity in EAE mice have been linked to neuroinflammation mediated by infiltrating T lymphocytes, activated microglia and reactive astrocytes (Centonze et al., 2009; Habbas et al., 2015; Mandolesi et al., 2013), we next examined the somatosensory cortex for the presence of inflammatory responses at acute disease. Gene expression analysis evidenced elevated levels of chemokines and adhesion molecules involved in the recruitment of peripheral immune cells such C-C motif chemokine ligand 2 (*Ccl2*), C-X-C motif chemokine ligand 10 (*Cxcl10*) and intercellular adhesion molecule 1 (*Icam*) (**Fig. 5a**). We also observed a significant increase of the microglia/activation marker cluster of differentiation 68 (*Cd68*) as well as elevated expression levels of pro-inflammatory factors and molecules involved in EAE and MS pathogenesis such as tumor necrosis factor *α* (*Tnfa*), interleukin 1α (*Il1a*), and components of complement system (**Fig. 5a**). In particular, the gene expression levels of *C1q* and *C1qb* were upregulated by 2-fold whereas complement component 3 (*C3*) was increased by > 200- fold in the EAE cortex. We next performed immunohistochemistry experiments using infiltrating and tissue resident cell markers to investigate the origin of cortical inflammation during EAE. Immunofluorescence staining of CD3 revealed the presence of scattered T cells associated to the meninges that correlated with neurological severity at acute EAE disease (Spearmańs r = 0.8710; *P* = 0.0023), but the cortex was not infiltrated (**Fig. 5c**). These results suggest that T cell infiltration is unlike to account for the cortical pathology associated to astrocyte dysfunction during EAE. We found elevated CD68 immunoreactivity and higher numbers of CD68-immunopositive pouches associated to cortical Iba-1 immunopositive profiles in cortical layers V-VI highlighting microglial activation in the somatosensory cortex of EAE mice at acute disease (**Fig. 5d-e**). Lastly, we examined the EAE cortex for the presence of astrocyte reactivity associated to aberrant calcium signaling during EAE. Immunofluorescence staining of the astrocyte reactivity marker GFAP evidenced increased expression levels in deep cortical layers from mice at acute disease that were corroborated by qPCR (**Fig. 5f-g** and **5b**). Previous studies have established that induction of astrocytic C3 is associated to the acquisition of disease phenotypes that promote neuronal and oligodendrocyte demise in EAE mice and human MS (Hou et al., 2020; Liddelow et al., 2017; Moreno-García et al., 2020). Hence, we next performed double immunolabeling using antibodies for complement component C3 and GFAP to decipher whether cortical astrocytes display increased levels of C3 during EAE. Consistent with gene expression results, GFAP and C3 immunoreactivity were increased in the cortical parenchyma of mice at acute EAE (**Fig. 5f-h**). Colocalization analysis showed that GFAP immunopositive astrocytic processes display increased C3 expression during EAE (**Fig. 5i**). Taken together, these results demonstrate that cortical autoimmune demyelination involves the emergence of reactive astrocytes in an inflammatory milieu that exhibit pathogenic potential.

### Neuroinflammatory mediators trigger aberrant astrocyte calcium signaling

We next asked whether astrocyte calcium signaling abnormalities in the cortex of EAE mice result from cell activation in an inflammatory milieu. To test this, we interrogated the calcium handling properties of astrocytes activated *in vitro* by incubation with the pro-inflammatory factors TNFα, IL-1α and C1q whose expression was significantly induced in the somatosensory cortex at acute disease (**Fig. 5a** and **f**). Astrocytes incubated with TNFα, IL-1α and C1q displayed upregulated expression of selected genes related to the phenotypic transformation of these cells during EAE and MS (Liddelow et al., 2017; Moreno-García et al., 2020) including complement component C3 (**Supplementary Fig. 7**). In parallel, we observed deregulated expression of a number of calcium signaling molecules including glutamate (*Grm1*, *Grm3*, *Grm5*) and ATP/adenosine receptors (*P2ry1*, *Adora1*), molecules related to endoplasmic reticulum calcium handling (*Itpr1*, *Ryr1*, *Ryr2*, *Ryr3*), and cytoplasmic Ca^2+^ binding proteins (*S100b*) (**Fig. 6a**). Some of these changes paralleled observations in astrocytes purified during acute EAE (**Fig. 4e**). Particularly, *Grm3* and *S100b* showed decreased gene expression in both experimental settings. However, genes such as *Grm1* and *Grm5* showed opposite changes in cells activated *in vitro* and *in vivo*.

**Figure 6.**
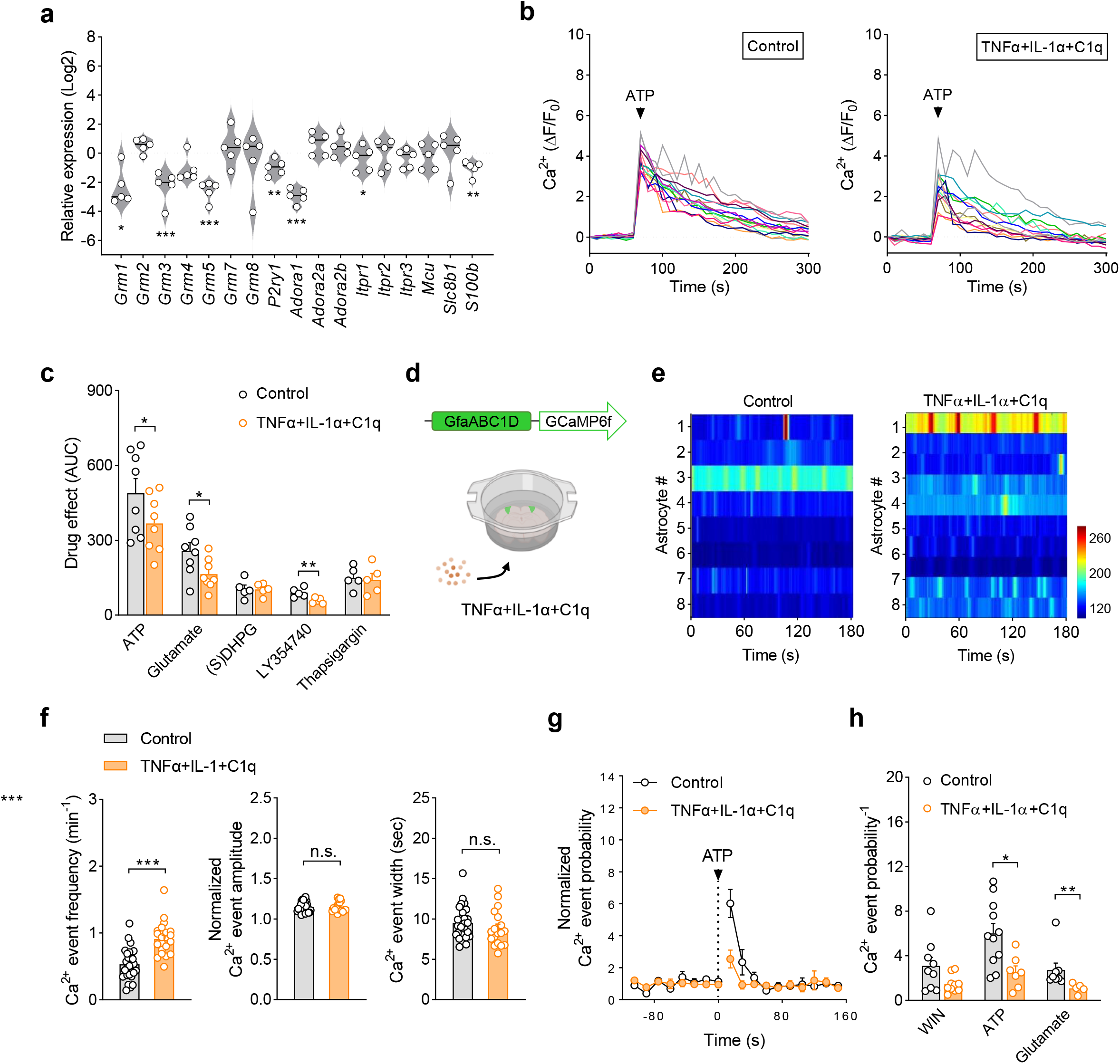
Pro-inflammatory signals engage aberrant astrocyte calcium activity. (**a**) Gene expression analysis of astrocyte calcium handling molecules in cultured cells activated *in vitro* by incubation with the pro-inflammatory factors TNFα (25 ng/ml), IL-1α (3 ng/ml) and C1q (400 ng/ml) (18-24 h). \**P* < 0.05, \*\**P* < 0.01; \*\*\**P* < 0.001; two-tailed paired Student’s *t* test or Mann Whitney test. (**b**) Representative imaging experiments showing calcium responses of individual astrocytes evoked by *in vitro* exposure to ATP (200 µM). (**c**) Quantitative analysis of calcium responses by ATP (200 µM), glutamate (200 µM), (S)DHPG (200 µM), LY354740 (200 µM), ATP (200 µM) and thapsigargin (1 µM) in astrocyte cultures incubated with TNFα+IL-1α+C1q. AUC, area under the curve. \**P* < 0.05, \*\**P* < 0.01; \*\*\**P* < 0.001; two-tailed paired Student’s *t* test or Mann Whitney test. (**d**) *Ex vivo* analysis of astrocyte calcium dynamics in response to pro-inflammatory factors. Acute cortical slices from mice injected with the AAV vector encoding for GCaMP6f under the Gfa-ABC1D promoter were imaged following incubation with TNFα+IL-1α+C1q (30 min). (**e**) Heat maps showing astrocytic calcium levels of individual astrocytes along time in cortical slices activated *ex vivo* with TNFα+IL-1α+C1q. (**f**) Frequency, amplitude and duration of spontaneous astrocyte calcium events in control conditions (28 slices, 5 mice) and following activation with pro-inflammatory factors (22 slices, 4 mice). \*\*\**P* < 0.001; two-tailed unpaired Student’s *t* test. (**g**) Calcium event probability normalized to basal activity over time depicting the effect of ATP (200 µM) in cortical astrocytes activated *ex vivo* with TNFα+IL-1α+C1q. (**h**) Calcium event probability normalized to basal activity depicting the effect of WIN55,212-2 (WIN; 300 µM), ATP (200 µM) or glutamate (200 µM) in control astrocytes (8-11 slices, 5 mice) and following incubation with pro-inflammatory factors (5-10 slices, 4 mice). \**P* < 0.05; \*\**P* < 0.01; two-tailed paired Student’s *t* test (before and after). n.s., not statistically significant. Error bars express SEM.

Next, we wondered whether the observed deficits in the expression calcium handling genes translated to deregulated cytosolic calcium dynamics in astrocytes activated *in vitro*. Calcium responses to the cannabinoid agonist WIN were below detection threshold in cultured astrocytes, probably reflecting low expression levels of the receptor protein in our *in vitro* experimental settings, but glutamate and ATP elicited reliable cytosolic calcium oscillations in control conditions (**Fig. 6b**, **c**). Cells activated by pro-inflammatory factors showed aberrant responses to glutamate, LY354740 and ATP and while the effect of (S)DHPG on cytosolic calcium levels remained unaffected (**Fig. 6c**). Incubation with the blocker of the sarco/endoplasmic reticulum calcium ATPase (SERCA) pump thapsigargin increased cytosolic calcium to the same extent in control and activated astrocytes, ruling out significant alterations in the calcium content of the ER (**Fig. 6c**). In order to corroborate these observations, we monitored astrocyte calcium responses using two-photon microscopy *ex vivo*. Cortical slices incubated with TNFα, IL-1α and C1q (30 min) showed increased calcium event probability of cortical astrocytes thus resembling astrocyte hyperactivity at acute EAE disease (**Fig. 6d-f**). However, the amplitude and duration of astrocytic calcium events in cortical slices incubated with the pro-inflammatory cocktail were similar to the control condition (**Fig 6f**). We next examined receptor-mediated astrocyte calcium responses in slices incubate with TNFα, IL-1α and C1. The calcium responses of cortical astrocytes to ATP and glutamate were significantly attenuated following short-term incubation with the pro-inflammatory cocktail (**Fig. 6g-h**). In parallel we observed a non-significant reduction in the ability of WIN to stimulate spontaneous astrocyte calcium activity (**Fig. h**). Thus, high local levels of pro-inflammatory mediators trigger deficits in the calcium responses of cortical astrocytes that mirror alterations during EAE pathology. Overall, these observations suggest that astrocyte activation in an inflammatory milieu underlie dysfunctional calcium dynamics with potential implication in neuronal network function.

### Cortical astrocyte-neuronal network interplay is disrupted during EAE

Calcium-based astrocyte signaling fine tunes neuronal activity through the release of gliotransmitters and deregulation of astrocyte calcium responses has the potential to contribute to cortical dysfunction in MS (Shigetomi et al., 2016). Hence, in order to decipher the physiopathological outcome of aberrant astrocyte calcium signaling on the activity of cortical networks we performed electrophysiological recordings of layer V pyramidal neurons. As first approach to investigate gliotransmission in the EAE cortex we monitored NMDA receptor-mediated slow inward currents (SICs) as electrophysiological readout for astrocytic glutamate release (Araque et al., 2014; Fellin et al., 2004; Perea & Araque, 2005). The amplitude (40.18 ± 5.74 pA EAE; 37.32 ± 3.21 pA control) and area (4216 ± 964 pA/ms EAE; 3998 ± 693 pA/ms control) of SICs in cortical layer V pyramidal neurons from immunized mice were similar to those recorded in control conditions. However, we found a significant increase in the frequency of SICs in EAE mice compared to naïve animals that resembles high levels of spontaneous astrocyte calcium activity measured *ex vivo* (**Fig. 7a-c**). The frequency of SICs was significantly reduced in IP_3_R_2_ ^-/-^ mice both in control conditions and during EAE thus corroborating the involvement of astrocyte calcium signaling in gliotransmitter release (**Fig. 7b**, **c**). Altogether, these results suggest that spontaneous calcium hyperactivity increases astrocytic glutamate release in the EAE cortex.

**Figure 7.**
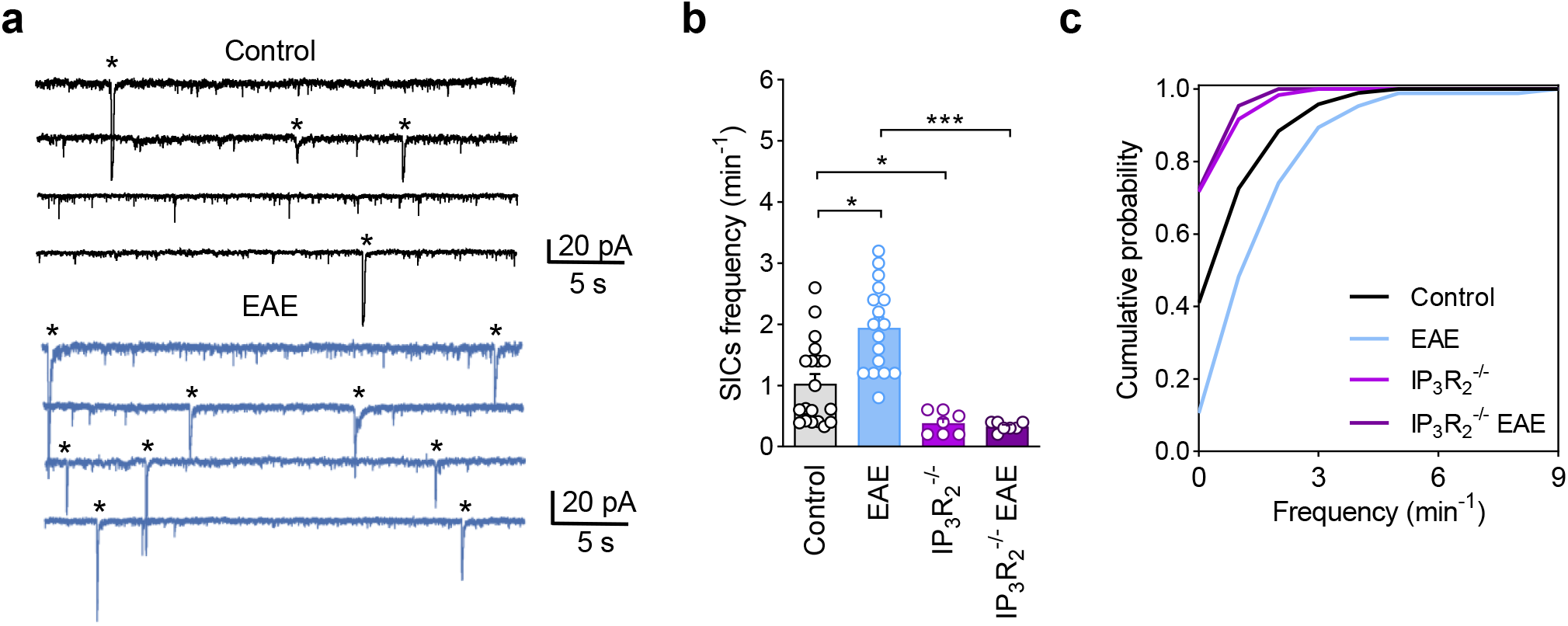
Exacerbated astrocytic glutamate release in the EAE cortex. (**a**) Representative traces depicting slow inward currents (SICs) (asterisks) in layer V pyramidal neurons within the somatosensory cortex from control and EAE mice. (**b**) Frequency of SICs in control (19 neurons, 4 mice), EAE (17 neurons, 6 mice), IP_3_R ^-/-^ (7 neurons, 2 mice) and IP_3_R_2_^-/-^ EAE (7 neurons, 2 mice) animals. \**P* < 0.05; \*\**P* < 0.01; one-way ANOVA, Dunn’s test for multiple comparisons. Error bars express SEM. (**c**) Cumulative frequency of SICs in cortical neurons from control, IP_3_R ^-/-^ and IP R ^-/-^ mice. Maximum differences between cumulative distributions were 0.304644 (EAE vs control), 0.306140 (IP_3_R_2_^-/-^ vs control) and 0.6171946 (IP_3_R ^-/-^ EAE vs EAE).

We next assessed cortical synaptic plasticity mechanisms that rely on astrocyte gliotransmitter release. We have recently shown that neuron-released endocannabinoids and astrocyte CB_1_Rs trigger short-term heteroneuronal synaptic depression mediated by ATP/adenosine as gliotransmitters in the somatosensory cortex (Baraibar et al., 2022). Thus, we next performed double patch-recordings of layer V pyramidal neurons and monitored excitatory postsynaptic currents (EPSCs) evoked by electrical stimulation of layer II-III in naïve and EAE mice at acute disease (**Fig 8a**). Consistent with our previous observations, depolarization of single layer V pyramidal neurons (ND) induced a transient depression of the synaptic transmission in 14 out of 42 (33%) heteroneurons (**Fig. 8b**, **d**) in non-immunized mice. We also observed a transient potentiation of the synaptic transmission in 10 out of 42 (24%) heteroneurons (**Fig. 8c**, **d**). Both heteroneuronal synaptic depression and potentiation were abolished following bath perfusion of the CB_1_ receptor antagonist AM251 (Baraibar et al., 2022) (**Fig. 8e**). EAE induction did not modulate the size of heteroneuronal synaptic plasticity (depression or potentiation) (**Fig. 8b**, **c**). However, the number of cortical heteroneurons exhibiting synaptic depression after ND was drastically reduced during EAE (4 out of 49 cells; 8%) (**Fig. 8d**). Conversely, immunized mice showed a significant increase in the percentage of transiently potentiated heteroneuronal synapses following ND (21 out of 49 cells; 43%) (**Fig. 8d**). Consistent with CB_1_ receptor mediated regulation, heterosynaptic depression and potentiation in EAE mice were prevented by bath application of AM251 (**Fig. 8e**). To further corroborate the role of astrocytic CB_1_ receptors in heterosynaptic plasticity during EAE we selectively deleted the *Cnr1* gene in cortical astrocytes by viral expression of Cre-recombinase under the control of the astroglial promoter GFAP in the somatosensory cortex of CB ^f/f^ mice (aCB ^-/-^) (Baraibar et al., 2022). Heteroneuronal depression was fully abolished in aCB_1_ ^-/-^ mice in both in control conditions (0 out of 13) and during EAE (0 out 15). In addition, heteroneuronal potentiation was also markedly reduced in naive (1 out of 13) and EAE (1 out of 15) mice. These observations suggest that EAE shifts short-term plasticity mediated by astrocyte CB_1_ receptors towards synaptic potentiation.

**Figure 8.**
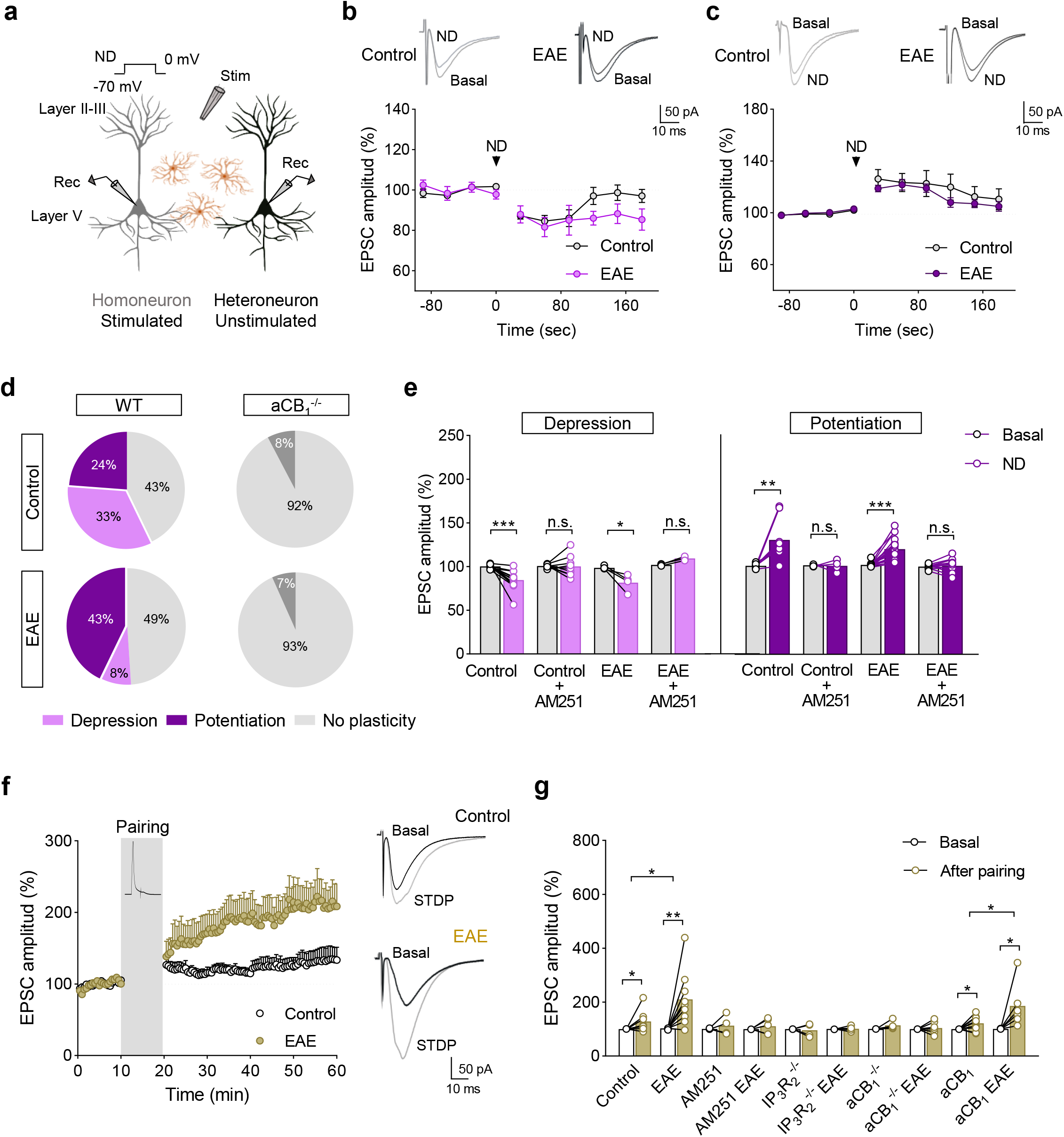
EAE disrupts astrocyte-mediated cortical neuron network plasticity. (**a**) Double patch-recordings from layer V pyramidal neurons in the somatosensory cortex for the analysis of astrocyte-neuron communication. (**b**-**c**) Time-courses and representative EPSC traces show heteroneuronal depression (**b**) and potentiation (**c**) of synaptic transmission following homoneuronal depolarization (ND) in control and EAE mice. (**d**) Percentage of heteroneurons that showed depression, potentiation or no plasticity in wild-type (WT) control (42 pairs, 7 mice) or EAE mice (49 pairs, 9 mice) and CB_1_^f/f^ mice injected with AAV-GFAP-Cre in the somatosensory cortex (aCB_1_^-/-^) in control conditions (13 pairs, 2 mice) and during EAE (15 pairs, 2 mice). \*\*\**P* < 0.001; Chi-square test. (**e**) Relative changes in EPSC amplitude depict heteroneuronal depression and potentiation in control (14 and 10 pairs, 6-5 mice), AM251 (2 µM) (9 and 4 pairs, 5-4 mice), EAE (4 and 21 pairs, 3-8 mice), EAE+AM251 (3 and 12 pairs, 2- 8 mice). \*\**P* < 0.01; \*\*\**P* < 0.001; two-tailed paired Student *t* test. (**f**) Time-course of EPSC amplitude in layer V pyramidal neurons depicts spike-timing dependent plasticity (STDP) induced in control and EAE mice by pairing a postsynaptically evoked action potential with an EPSP (Δt = −25 ms) 60 times in 10 minutes (gray area). Error bars express SEM. (**g**) Relative changes in EPSC amplitude after the pairing period in control and immunized mice (8 and 10 neurons, 4-7 mice), in the presence of AM251 (4 and 5 neurons, 2-3 mice), in IP_3_R_2_^-/-^ animals (4 and 4 neurons, 2-2 mice), and in aCB_1_R^-/-^ (5-7 neurons, 2-3 mice) or aCB_1_R (9 and 6 neurons, 3-3 mice) mice injected with AAV-GFAP-Cre and AAV-GFAP-mCherry, respectively. \**P* < 0.05; \*\**P* < 0.01; two-tailed paired Student’s *t* test (before and after) and unpaired t tests (between groups). n.s., not statistically significant. Error bars express SEM.

To further corroborate the hypothesis that autoimmune inflammation impairs astrocyte-mediated modulation of cortical synaptic function we next investigated long-term plasticity mechanisms that depend on astrocytic calcium signaling during EAE. Endocannabinoids and astrocyte CB_1_ receptors gate a form of spike timing-dependent long-term plasticity (STDP) in the neonatal mouse that relies on the astrocytic delivery of glutamate as gliotransmitter and switches to long-term potentiation at more mature stages (Martínez-Gallego et al., 2022; Min & Nevian, 2012). Thus, we next investigated the possible alterations in STDP in the EAE cortex. With this aim, we recorded EPSCs at layer II-III to layer V synapses onto pyramidal neurons and induced STDP by pairing each EPSC with a preceding postsynaptically evoked action potential (AP-EPSP pairing, timing interval Δt = −25 ms, 60 AP-EPSP pairings in 10 minutes). This stimulation protocol induced mild long-term potentiation of synaptic transmission in good agreement with previous observations in the adult mouse neocortex (**Fig. 8f-g**) (Martínez-Gallego et al., 2022). Remarkably, the STDP induction protocol resulted in a significantly augmented potentiation of cortical synaptic transmission in EAE mice (**Fig. 8f-g**). We next aimed at corroborating the role of astrocyte CB_1_Rs as mediators of STDP in control and disease conditions. Bath application of AM251 prevented the expression of STDP both in control conditions and during EAE suggesting that both phenomena were mediated by CB_1_ receptors (**Fig. 8g**). STDP was also abolished in control and immunized aCB_1_ ^-/-^ mice further suggesting that astrocyte CB_1_ receptor trigger the potentiation of excitatory transmission in naïve and immunized mice (**Fig. 8g**). The STDP induction protocol elicited a slight potentiation of synaptic transmission in CB_1_^f/f^ mice injected with AAV8-GFAP-mCherry (aCB_1_) that was significantly augmented during acute EAE thus corroborating the results obtained in non-transgenic mice (**Fig. 8g**). Finally, we examined the dependency of STDP on astrocytic calcium in control and disease conditions using IP_3_R_2_ ^−/−^ mice in which G protein-mediated Ca^2+^ signal is selectively impaired in astrocytes (Baraibar et al., 2022; Di Castro et al., 2011). Consistent with our observations in aCB_1_ ^-/-^ mice, the spike-timing dependent forms of potentiation recorded in control conditions and at acute EAE disease were absent in IP_3_R_2_ ^-/-^ mice (**Fig. 8g**). Altogether, these results show that EAE disrupts cortical neuron network plasticity mechanisms triggered by astrocyte CB_1_ receptors causing neuronal network hyperactivity.

## Discussion

Cortical dysfunction is a core pathological feature of MS that underlies cognitive deficits and predicts disease progression (Calabrese et al., 2015; Mahad et al., 2015). Neuronal and synaptic failure have been studied extensively in patients and animal models of the disease and are postulated at the origin of functional deficits in cortical MS (Jafari et al., 2021; Jürgens et al., 2016). Astrocytes fine-tune neuronal network operation through the release of gliotransmitters (Araque et al., 2014) and have been crucially involved in MS pathophysiology (Linnerbauer et al., 2020). However, the contribution of astroglial cells to MS cortical network dysfunction has remained largely unknown. In this study we investigated the calcium activity of cortical astrocytes and the features of gliotransmission in a mouse model of MS. Applying fiber photometry calcium imaging in freely-behaving mice we discovered the emergence of dysfunctional spontaneous and stimulus-evoked astrocyte activity patterns associated to EAE progression. Cortical astrocytes in EAE mice showed spontaneous calcium transients of reduced amplitude and frequency but increased duration, whereas sensory-evoked calcium events were impaired only in terms of amplitude. The onset of dysfunctional astrocytic calcium activity paralleled the appearance of neurological disability thus raising the possibility that these changes may reflect motor deficits. Indeed, locomotion is directly associated to increases in astrocytic calcium activity *in vivo* (Bojarskaite et al., 2020; Qin et al., 2020) and it seems plausible that the loss of motor coordination and muscle tone associated to EAE progression underlies, at least in part, aberrant astrocyte calcium dynamics in the somatosensory cortex. Supporting this hypothesis, deficits in the amplitude of astrocyte calcium transients correlated to disease severity at different time-points of disease progression. By contrast, neither the increase in the duration of astrocyte calcium events nor the frequency deficits correlated with motor symptomatology during EAE, which suggests that different disease-associated regulatory mechanisms mediate changes in each parameter. In this regard, our two-photon imaging analysis of astrocyte calcium dynamics in cortical slices unveiled a hyperactive phenotype in terms of event probability, amplitude and duration, in line with emerging studies on astrocyte dysfunction in neurodegenerative and neuroinflammatory conditions such as Alzheimeŕs disease (Delekate et al., 2014; Kuchibhotla et al., 2009; Lines et al., 2022; Shah et al., 2022), Parkinsońs disease (Nanclares et al., 2023).

Astrocyte calcium hyperexcitability in the EAE cortex was encompassed by diminished responses mediated by astroglial CB_1_ receptors both *in vivo* and *ex vivo*. This functional deficit might reflect reductions in the astrocyte-associated receptor pools during acute disease (Moreno-García et al., 2020) and/or be associated to the deregulation of intracellular calcium signaling machinery as suggested by our targeted gene expression analysis. Notably, the astrocyte hypo-responsiveness to CB_1_ receptor activation was mirrored by deficits in the calcium activity of astrocytes to ATP, glutamate and selective mGluR_1/5_ and mGluR_2/3_ agonists further suggesting a widespread deregulation of astrocyte calcium signaling pathways during autoimmune inflammation. These observations are reminiscent of recent reports of spontaneous hyperactivity but reduced sensory-evoked astrocyte responsiveness in experimental Alzheimeŕs disease (Lines et al., 2022). Irrespective of the precise mechanisms underlying astrocyte calcium hypo-responsiveness to acute activation, the emerging picture is that impaired astrocyte activity mediated by endocannabinoids, ATP/adenosine or glutamate as neurotransmitters may contribute to dysfunctional spontaneous and sensory-evoked calcium oscillations in the MS cortex.

Activation of astrocytes by either G_i_- or G_q_-DREADDs increased the frequency and amplitude of calcium signals as consistent with previous studies using pharmacogenetic tools (Baraibar et al., 2022; Durkee et al., 2019), as well as pharmacological or sensory-evoked astrocyte stimulation (Ding et al., 2013; Jiang et al., 2016; Lines et al., 2020; Wang et al., 2006). Mechanistically, we show that deficits in astrocyte calcium responses to neurotransmitter receptors in the EAE cortex rely on impaired activity of both G_i_ and G_q_ protein signaling pathways as concluded from the impaired responses to pharmacological activation of DREADDs expressing cells. An interesting observation from our pharmacogenetic experiments is that signaling mediated by astrocyte G_i_ proteins is predominantly altered in cortical astrocytes during autoimmune inflammation as compared to G_q_ calcium signaling. These observations confirm the importance of the G_i_ protein signaling pathway in astrocyte physiology and highlight its relevance for the acquisition of astrocyte disease phenotypes linked to MS.

In this study we establish a mechanistic association between cortical autoimmune neuroinflammation and the emergence of aberrant astrocyte phenotypes at the calcium excitability level. The presence of reactive astrocytes expressing complement C3 was linked to elevated levels pro-inflammatory factors in the EAE cortex including the microglial-secreted cytokines TNFα, IL-1α and C1q (Clarke et al., 2018; Liddelow et al., 2017). Remarkably, acute slice activation with a pro-inflammatory cocktail composed of TNFα, IL-1α and C1q reproduced spontaneous astrocyte hyperactivity at the calcium event frequency level and hypo-responsiveness to ATP and glutamate thus mirroring observations in the EAE cortex. On mechanistic grounds, using an *in vitro* setting we demonstrate cell-autonomous effects of pro-inflammatory astrocyte activation with TNFα, IL-1α and C1q on the expression of calcium handling molecules encompassed by dysfunctional calcium responses. Altogether these results highlight the relevance of TNFα, IL-1α and C1q as major upstream mediators of dysfunctional astrocyte calcium signaling in the MS cortex. Future assessment in triple knockout mice lacking expression of these cytokines (Clarke et al., 2018; Hartmann et al., 2019) may help elucidate the relevance of these pro-inflammatory signals in the emergence of aberrant astrocyte signaling and their potential for astrocyte therapeutic targeting in cortical MS.

Our observations in cultured cells are in good agreement with previously published results showing that inflammatory astrocyte activation with TGF-ß1, LPS and IFNγ significantly deregulates, mostly down, astrocyte calcium signaling both at the gene expression and functional level in parallel to the upregulation of immune signaling and cell injury molecular networks (Hamby et al., 2012). Our gene expression results in cultured cells highlight commonly down-regulated genes, such as *P2ry1* and *Adora1*, but also certain discrepancies most likely related to the different pro-inflammatory stimuli used. In the same line, we observed several inconsistencies regarding deregulation of calcium handling molecules between EAE astrocytes and cultured cells activated with TNFα, IL-1α and C1q which may reflect a more complex and dynamic deregulation of calcium signaling in astroglia activated in the context of autoimmune demyelination. Altogether these results support the emerging concept that reactive astrogliosis is associated to neuroinflammatory context-specific deficits in calcium signaling mechanisms that feature heterogeneous populations of dysfunctional astrocytes contributing to neuropathologies (Escartin et al., 2021; Hasel et al., 2021).

A remarkable finding in this study is that altered astrocyte calcium excitability encompasses aberrant cortical gliotransmission during autoimmune inflammation. We show that the frequency of SICs is increased in the EAE cortex highlighting an exacerbated release of glutamate from reactive astrocytes that mirrors recent observations in rodent models of Alzheimeŕs disease (Lines et al., 2022; Talantova et al., 2013). Furthermore, our results unveil that short-term and long-term plasticity mechanisms mediated by astrocyte CB_1_ receptors in the brain cortex (Baraibar et al., 2022; Martínez-Gallego et al., 2022) shift to potentiation during autoimmune demyelination. Thus, deficits in astrocyte CB_1_ receptor mediated calcium signaling translate into gliotransmission defects leading to the potentiation of cortical excitation. Strikingly, our electrophysiological results suggest that heterosynaptic and spike-timing dependent plasticity mechanisms in the inflamed cortex still depend on intracellular calcium mobilization following the activation of the astrocytic CB_1_ receptor pool. Altogether, our results suggest that functional reshaping of astrocyte-neuron networks during autoimmune demyelination involves complex adaptations in calcium signal-to-noise ratio that lead to deregulation in the identity and/or relative amount of gliotransmitters shaping cortical synaptic excitation.

Concerning the pathogenic implications of dysfunctional astrocyte calcium signaling in cortical MS emerging evidence shows that deregulation in the calcium dynamics of astroglial cells affect neuronal network function leading to behavioral abnormalities in neurodegenerative diseases (Nagai et al., 2021; Shah et al., 2022; Shigetomi et al., 2016). In this context, the results of the present study suggest that dysfunction of astrocyte-neuronal interplay may deregulate cortical electrical activity and contribute to cognitive decline in MS patients. Also related to the pathogenesis of cognitive and memory impairments, it is well established that MS cortical pathology involves early excitatory-inhibitory imbalance leading and synaptic loss and neurodegeneration (Calabrese et al., 2015; Ellwardt et al., 2018; Jafari et al., 2021; Jürgens et al., 2016; Potter et al., 2016). Of note, exacerbated release of glutamate from astrocytes is postulated to contribute to neuronal excitotoxicity and synapse loss in status epilepticus and Alzheimeŕs disease supporting the neurotoxic potential of aberrant astrocytic gliotransmission in neurodegenerative and neuroinflammatory conditions (Ding et al., 2007; Talantova et al., 2013). In this study, dysfunctional astrocyte calcium activity and gliotransmission were not associated to significant synaptic deficits or synapse loss in the EAE cortex. However, our electrophysiological analysis shows increased duration of mEPSCs in layer V pyramidal neurons reminiscent of previous observations in the striatum and cerebellum of EAE mice (Centonze et al., 2009; Mandolesi et al., 2013). These observations suggest that aberrant gliotransmission encompasses early excitatory synaptic imbalance in the MS cortex and supports the possibility that astrocyte dysfunction leading to exacerbated excitatory transmission contributes to synaptic deregulation and neurodegeneration in cortical MS.

In summary, the present study provides key findings on the role of astrocytes in cortical MS pathology. We have shown that cortical astrocytes activated in an inflammatory milieu are spontaneously hyperactive but hypo-responsive to sensory-stimulus evoked and receptor-mediated calcium signaling. Aberrant calcium excitability encompasses astrocyte-to-neuron communication defects leading to the potentiation of excitatory transmission. These observations suggest that aberrant astrocyte calcium signaling and gliotransmission contribute to abnormal cortical network operation in MS thus pointing to novel therapeutic targets against cognitive decline.

## Supporting information

Supplementary Video 1. Basal Control

Supplementary Video 2. Basal EAE

Supplementary Video 3. Duration Control

Supplementary Video 4. Duration EAE

Supplementary Video 5. WIN Puff Control

Supplementary Video 6. WIN Puff EAE

---

**Supplementary Figure 1.**
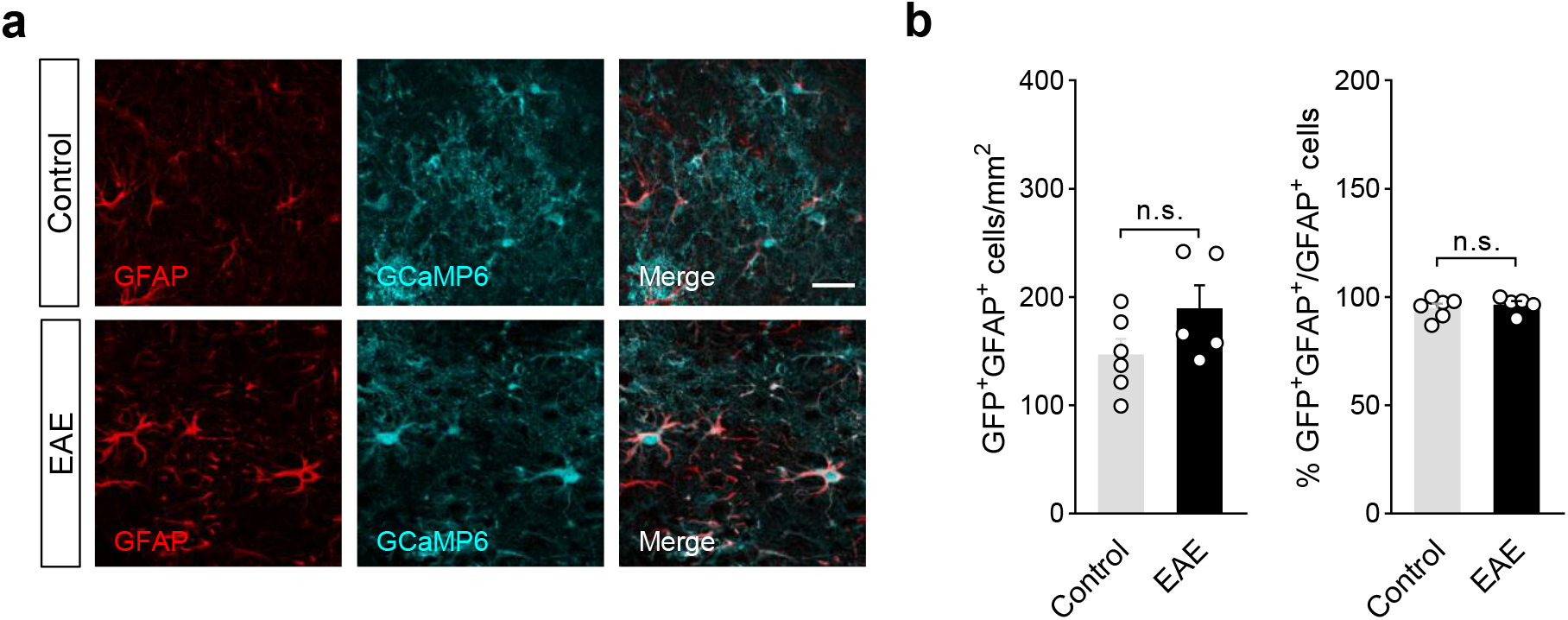
Imaging astrocyte calcium activity in the somatosensory cortex of EAE mice. (**a**) Representative images of control and EAE mouse cortices injected with AAV-GFAP-GCaMP6s and stained for GFP (green) and for the astrocyte marker GFAP (red). Scale bar: 25 µm. (**b**) Quantitative analysis of GFAP and GFP colocalization in control (6 mice) and immunized animals (21 dpi; 5 mice). Data were analyzed by two-tailed unpaired Student *t* test. n.s., not statistically significant. Error bars express SEM.

**Supplementary Figure 2.**
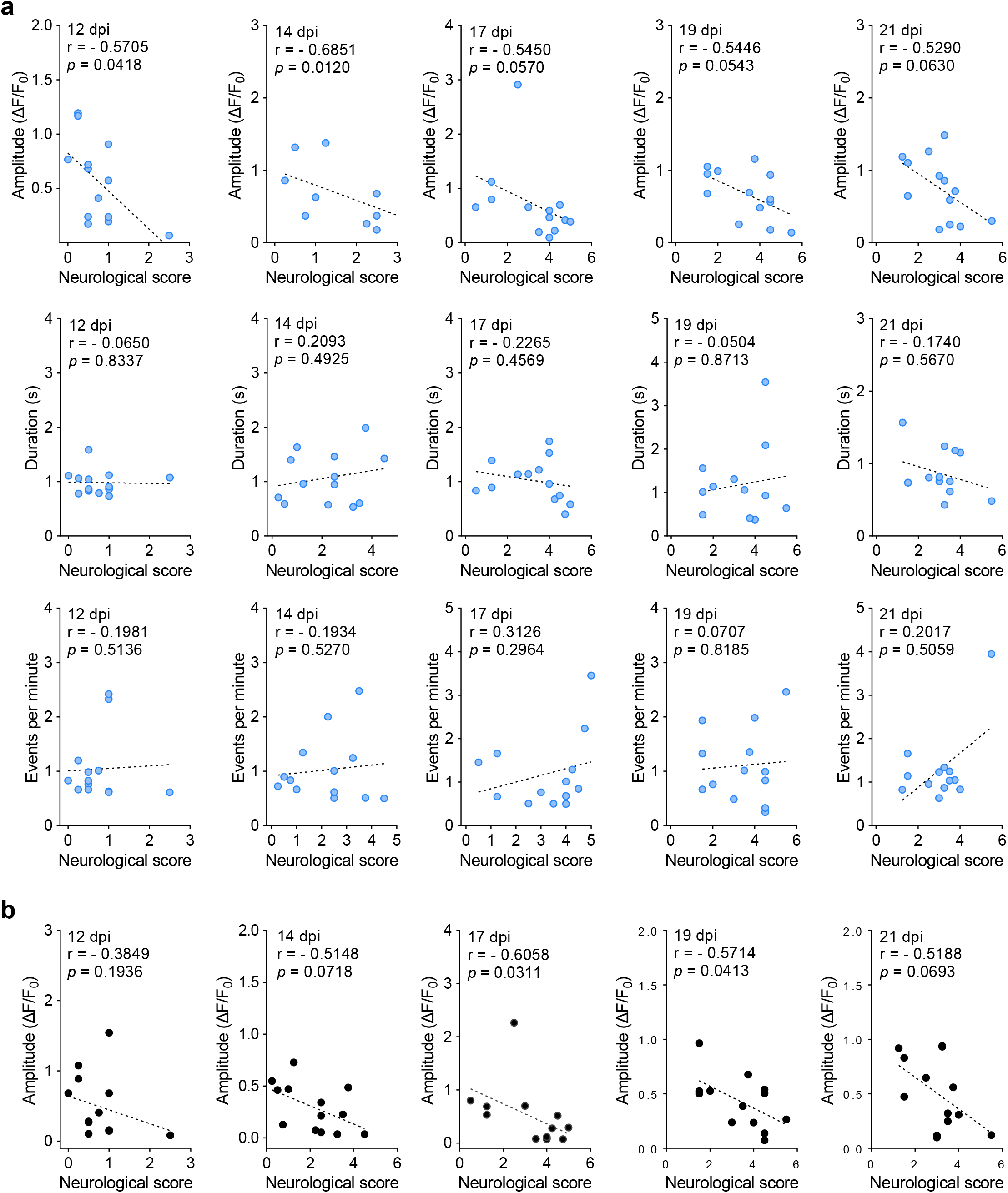
Dysregulation of astrocyte calcium activity in the EAE cortex: correlation to disease severity. (**a**) Correlation analysis between the amplitude (top), duration (middle) and frequency (bottom) of spontaneous astrocyte calcium oscillations in the mouse cortex and neurological score at different time-points of EAE disease progression (13 mice). (**b**) Correlation analysis between the calcium responses evoked by tail-holding in cortical astrocytes during EAE. Raw data from EAE mice were normalized to values recorded during the pre-symptomatic phase and plotted against neurological scores. Pearson’s or Spearman’s correlation coefficients and *P* values are indicated in each dot plot.

**Supplementary Figure 3.**
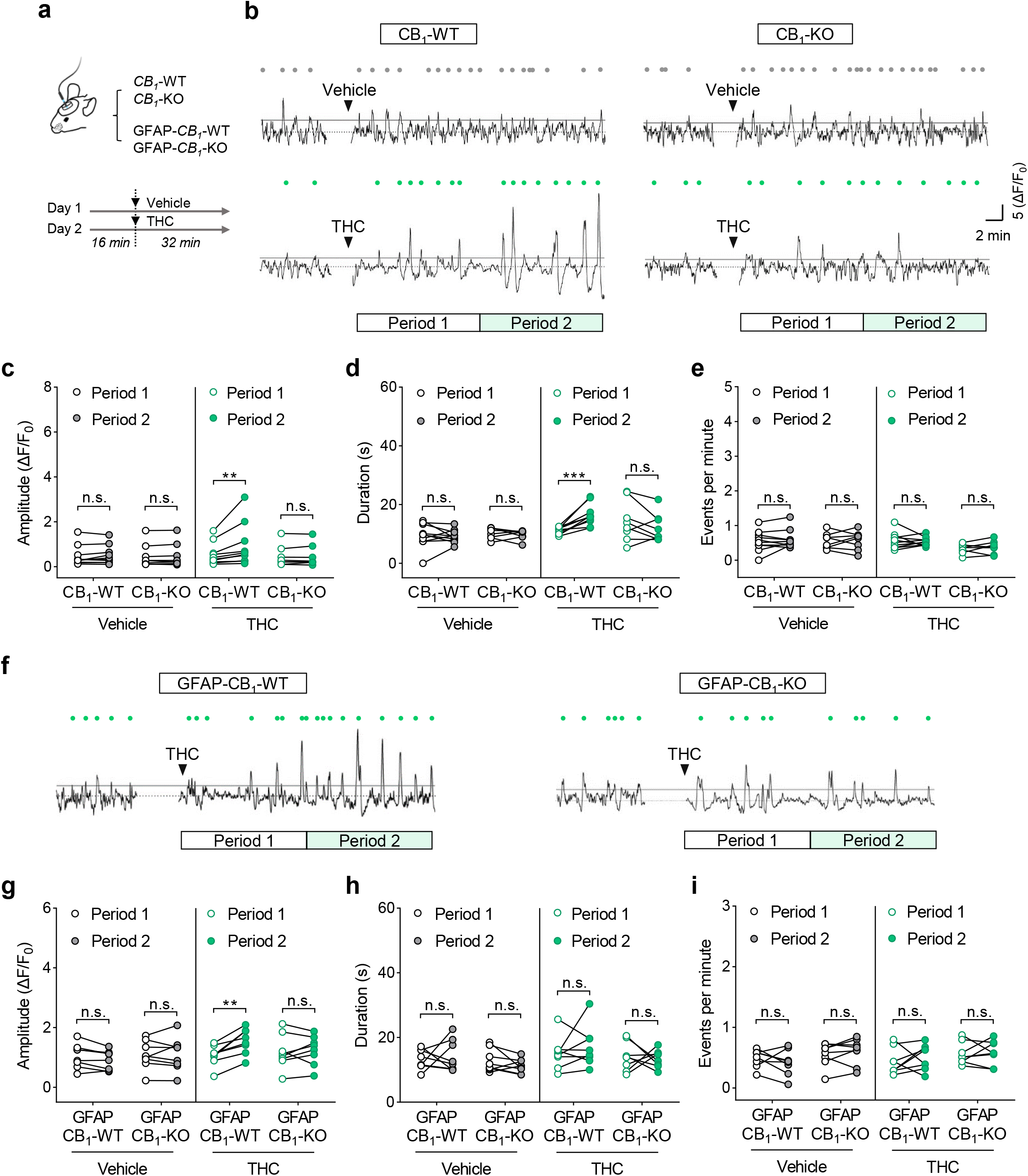
Astrocyte CB_1_ receptors induce cytosolic calcium oscillations *in vivo*. (**a**) Experimental fiber photometry approach for *in vivo* imaging of astrocyte calcium responses to THC in transgenic mice. (**b**) Representative traces from the somatosensory cortex of CB_1_-WT (left) and CB_1_-KO (right) mice injected with AAV-GFAP-GCaMP6fs and treated with vehicle (top) and THC (10 mg/Kg; i.p.) (bottom). (**c**- **e**) Comparison of amplitude (**c**), duration (**d**) and frequency (**e**) of calcium oscillations recorded from CB_1_-WT (10 mice) and CB_1_-KO (8 mice) animals following vehicle or THC injection. (**f**) Representative recordings from the somatosensory cortex of GFAP-CB_1_-WT (left) and GFAP-CB_1_-KO (right) mice injected with AAV-GFAP-GCaMP6fs and treated with THC. (**g**-**i**) Amplitude (**g**), duration (**h**) and frequency (**i**) of calcium transients in GFAP-CB_1_-KO mice (8 mice) and GFAP-CB_1_-WT control littermates (8 mice) following vehicle and THC injection. Gray and green dots in **b** and **f** correspond to detected transients above the threshold (median+2*MAD) in mice injected with vehicle and THC. White and green rectangles show the time window of analyzed period 1 and period 2, respectively. The first minute before and after injection was removed to exclude mouse-handling/injection effects. \*\**P* < 0.01; \*\*\**P* < 0.001; two-way ANOVA followed by Šídák’s multiple comparisons test. n.s., not statistically significant.

**Supplementary Figure 4.**
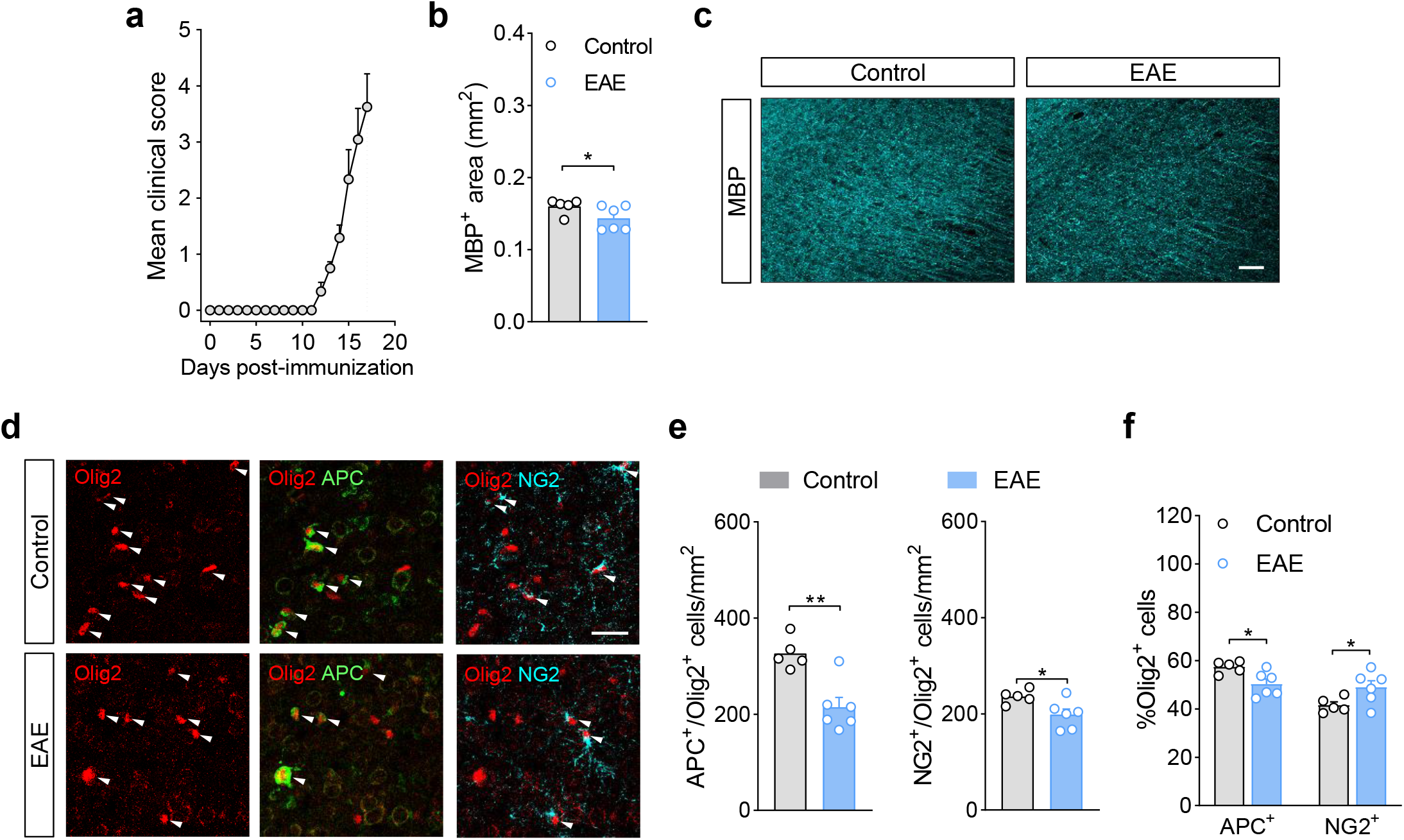
Cortical demyelination and oligodendrocyte loss at acute EAE disease. (**a**) Time-course of neurological disability in EAE mice used for immunohistochemical analysis of the somatosensory cortex at acute disease (17 dpi). (**b**) Quantification of MBP in immunostained cortical layer V-VI for control (5 mice) and EAE animals (6 mice). \**P* < 0.05; two-tailed unpaired Student´s *t* test. (**c**) Representative confocal images for MBP immunolabeling in layer V-VI of the somatosensory cortex in control and EAE conditions. Scale bar = 50 µM. (**d**) Representative labelling of oligodendrocyte markers Olig2, APC and NG2 in layer VI of the somatosensory cortex. Scale bar = 25 µM. (**e**) Quantification of Olig2/APC-positive mature oligodendrocytes (left) and Olig2/NG2-positive oligodendrocyte progenitor cells (right) in cortical layer V-VI for control and EAE mice. \**P* < 0.05; \*\**P* < 0.01; two-tailed unpaired Student´s *t* test. (**f**) Percentages of oligodendrocytes (APC^+^) and progenitor cells (NG2^+^) in the Olig2-positive population indicate a reduced proportion of mature cells in the EAE cortex. \**P* < 0.05; two-tailed unpaired Student´s *t* test.

**Supplementary Figure 5.**
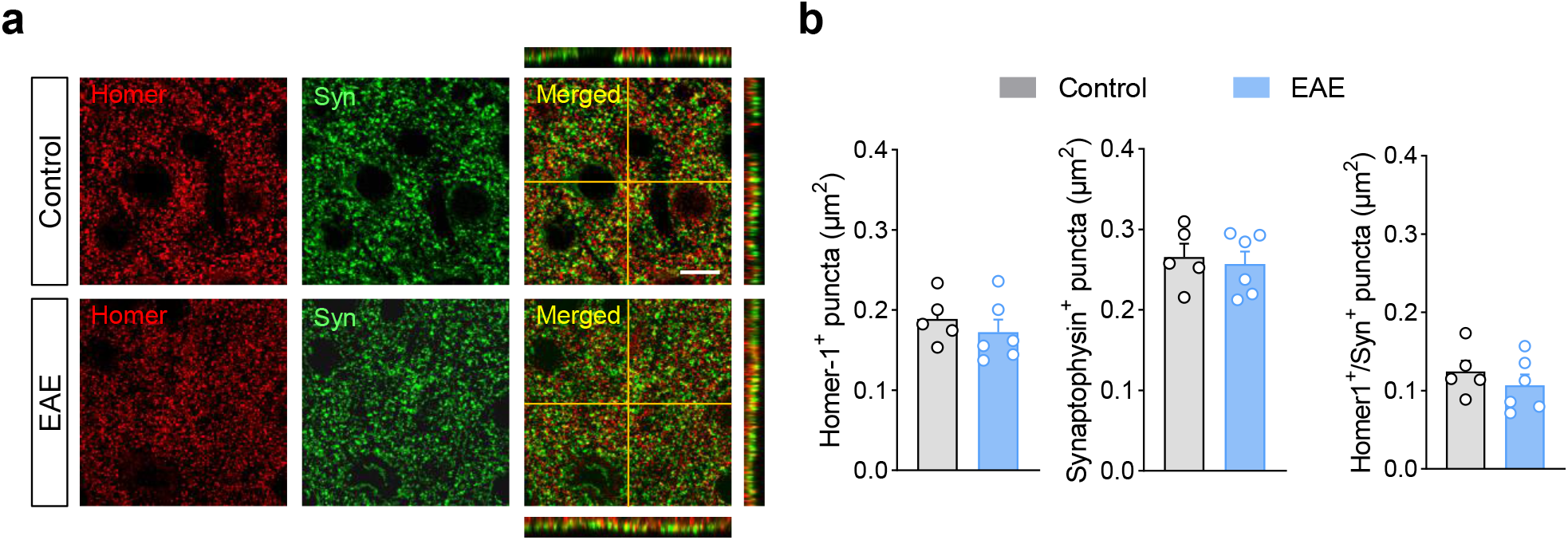
Intact excitatory synaptic inputs in the EAE cortex. (**a**) Representative labeling of excitatory presynaptic (Synaptophysin, Syn) and postsynaptic (Homer-1) markers in somatosensory cortex layer V. Scale bar = 10 µM. (**a**) Quantification of cortical excitatory synapse density in control (5 mice) and EAE conditions (6 mice). Error bars express SEM.

**Supplementary Figure 6.**
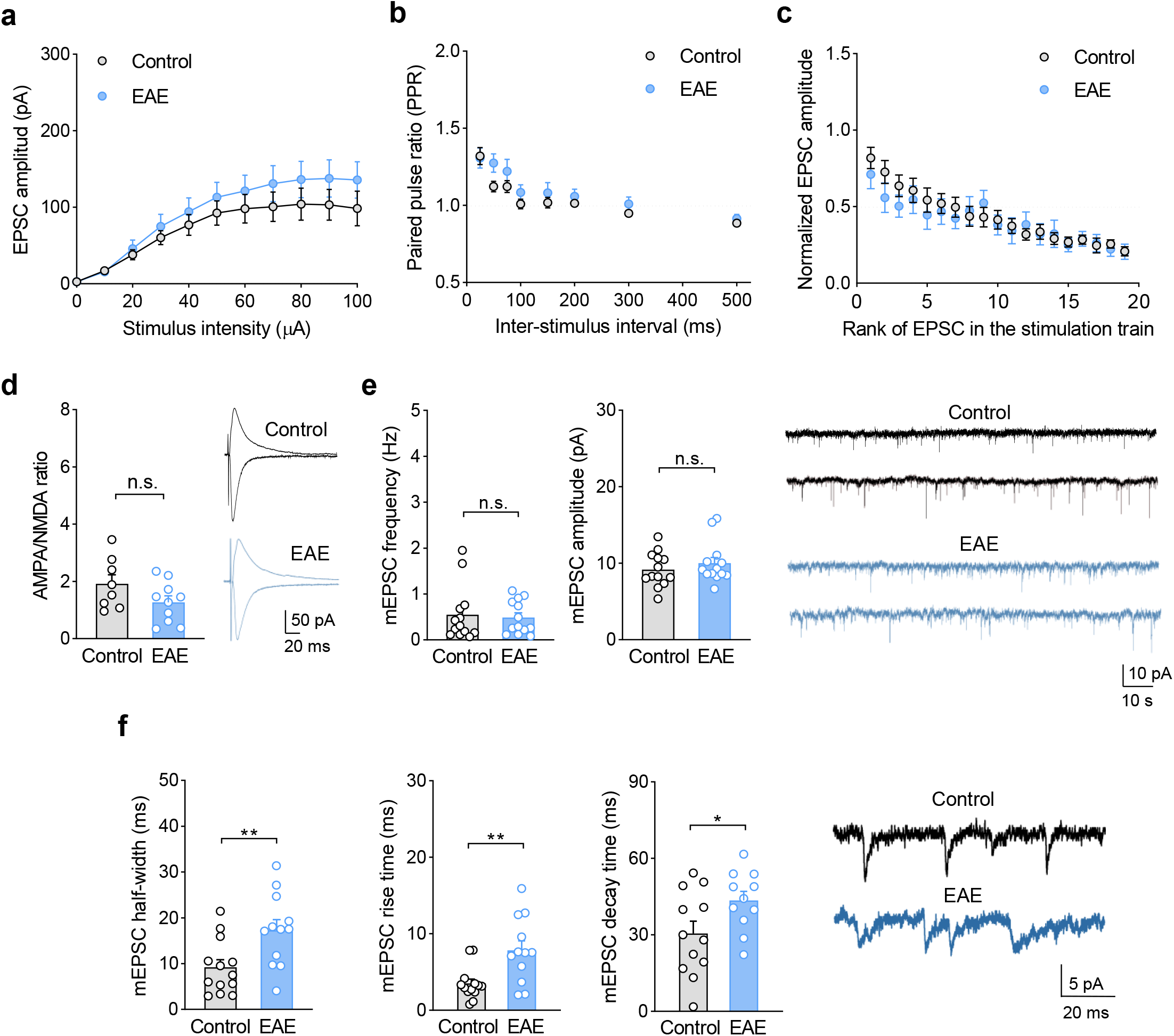
Mild functional synaptic defects in the EAE cortex. (**a**) Input-output relationships of evoked EPSCs in somatosensory cortex layer V pyramidal neurons from control (11 neurons, 4 mice) and EAE (18 neurons, 5 mice) animals. (**b**) Paired-pulse ratio induced by two consecutive stimuli delivered at different time intervals in control conditions (14 neurons, 4 mice) and during EAE (13 neurons, 4 mice). (**c**) Synaptic fatigue induced by 19 consecutive stimuli at 25 ms interpulse intervals in control (15 neurons; 4 mice) and EAE (10 neurons, 4 mice) mice. (**d**) AMPA to NMDA receptor current ratio and representative traces from naïve animals (15 neurons, 2 mice) and mice and acute EAE disease (10 neurons, 3 mice). (**e**) Representative traces and quantitative analysis of mEPSC frequency and amplitude in control conditions (8 neurons, 4 mice) and during EAE (10 neurons, 6 mice). (**f**) Half-width, rise time, decay time and representative traces depict increased duration of mEPSCs in layer V neurons from EAE mice as compared to naïve animals. \**P* < 0.05; \*\**P* < 0.01; two-tailed unpaired Student’s *t* test. n.s., not statistically significant. Error bars express SEM.

**Supplementary Figure 7.**
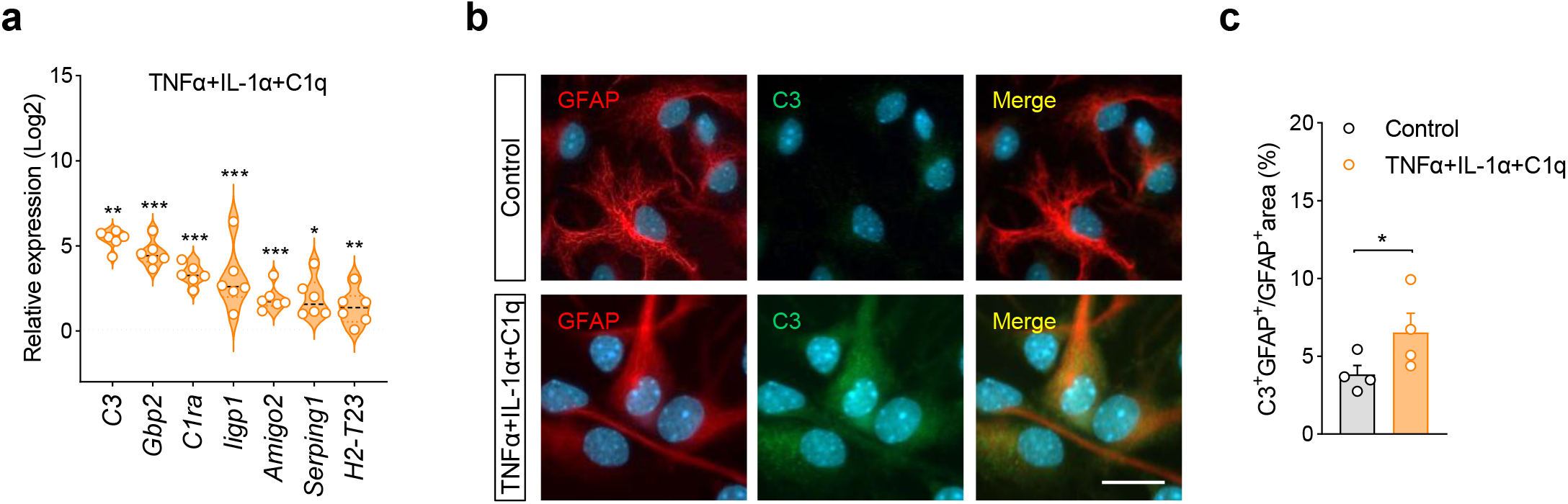
Pro-inflammatory activation of astrocytes *in vitro*. (**a**) Gene expression of neurotoxicity related molecules in astrocyte cultures activated with TNFα (25 ng/ml), IL-1α (3 ng/ml) and C1q (400 ng/ml) (18-24 h) relative to control cells (6 cultures). (**b**) Representative confocal images depict C3 immunolabelling in astrocyte cultures stimulated with pro-inflammatory factors. Scale bar = 25 µm. (**c**) Quantitative analysis of C3 expression in astrocytes *in vitro* (4 cultures). \**P* < 0.05; \*\**P* < 0.01; \*\*\**P* < 0.001; two-tailed paired Student´s *t* test. Error bars express SEM.

**Supplementary video S1.** Spontaneous calcium activity of cortical astrocytes in control conditions.

**Supplementary video S2.** Spontaneous calcium activity of cortical astrocytes during EAE (14 dpi).

**Supplementary video S3.** Representative astrocyte calcium event in a control mouse.

**Supplementary video S4.** Representative astrocyte calcium event during EAE (18 dpi).

**Supplementary video S5.** Calcium response of cortical astrocytes to the CB_1_ receptor agonist WIN55,212-2 in control conditions.

**Supplementary video S6.** Calcium response of cortical astrocytes to the CB_1_ receptor agonist WIN55,212-2 during EAE (14 dpi).

## Acknowledgements

We would like to thank the personnel of the Animal Facilities of the University of the Basque Country and Neurocentre Magendie for mouse care. We also thank G. Perea for providing IP_3_R_2_ ^-/-^ mice, S. Calovi and F.N. Soria and for support in the analysis of confocal images. This work was funded by FEDER and Instituto de Salud Carlos III (PI21/00629, to S.M. and A.R.-A; CB06/05/00, to C.M.), Basque Government (PIBA19- 0059 and IT1473-22, to S.M.; IT1203-19, to C.M.), ARSEP Foundation (to S.M. and G.M.), INSERM (to G.M.), the European Research Council (MiCaBra, ERC-2017-AdG- 786467, to G.M.), Fondation pour la Recherche Medicale (FRM, DRM20101220445 to G.M.), the Human Frontiers Science Program (to G.M.), Region Aquitaine (CanBrain, AAP2022A-2021-16763610 and -17219710 to G.M.); French State/Agence Nationale de la Recherche (CaCoVi, ANR 18-CE16-0001-02; MitObesity, ANR 18-CE14-0029- 01; ERA-Net Neuron CanShank, ANR-21-NEU2-0001-04, to G.M), Spanish Ministry of Science and Innovation (PID2019-109724RB-100 to C.M.; PGC2018-093990-A-I00, to E.S.-G.), National Institutes of Health-MH (MH, R01MH119355; NINDS, R01NS097312; NIDA, R01DA048822, to A.A.), Postdoctoral and Predoctoral Programs of the Basque Government (to A.M.-B., T.C. A.M.G., and C.U), Predoctoral Program of the UPV/EHU (to E.S.).

## Author contributions

G.M and S.M. conceptualized and supervised the study. A.M.B performed *ex vivo* electrophysiology and imaging experiments. T.C. conducted imaging and expression analyses in astrocyte cultures and purified cells. A.M.-G. and R.S. performed and supervised *in vivo* imaging experiments. T.C., E.S., C.L.U, A.B.-C. conducted and supervised mouse perfusion and immunohistochemistry experiments. F.S.-L. provided guidance for the analysis of confocal microscopy images. E.S.-G. provided some CB_1_^f/f^ mice to the group of S.M. A.R.-A. provided conceptual ideas. A.A. and C.M. provided data and conceptual support. A.M.B, G.M. and S.M. produced the figures and wrote the paper with input from all authors.

---

**Supplementary Table 1.**
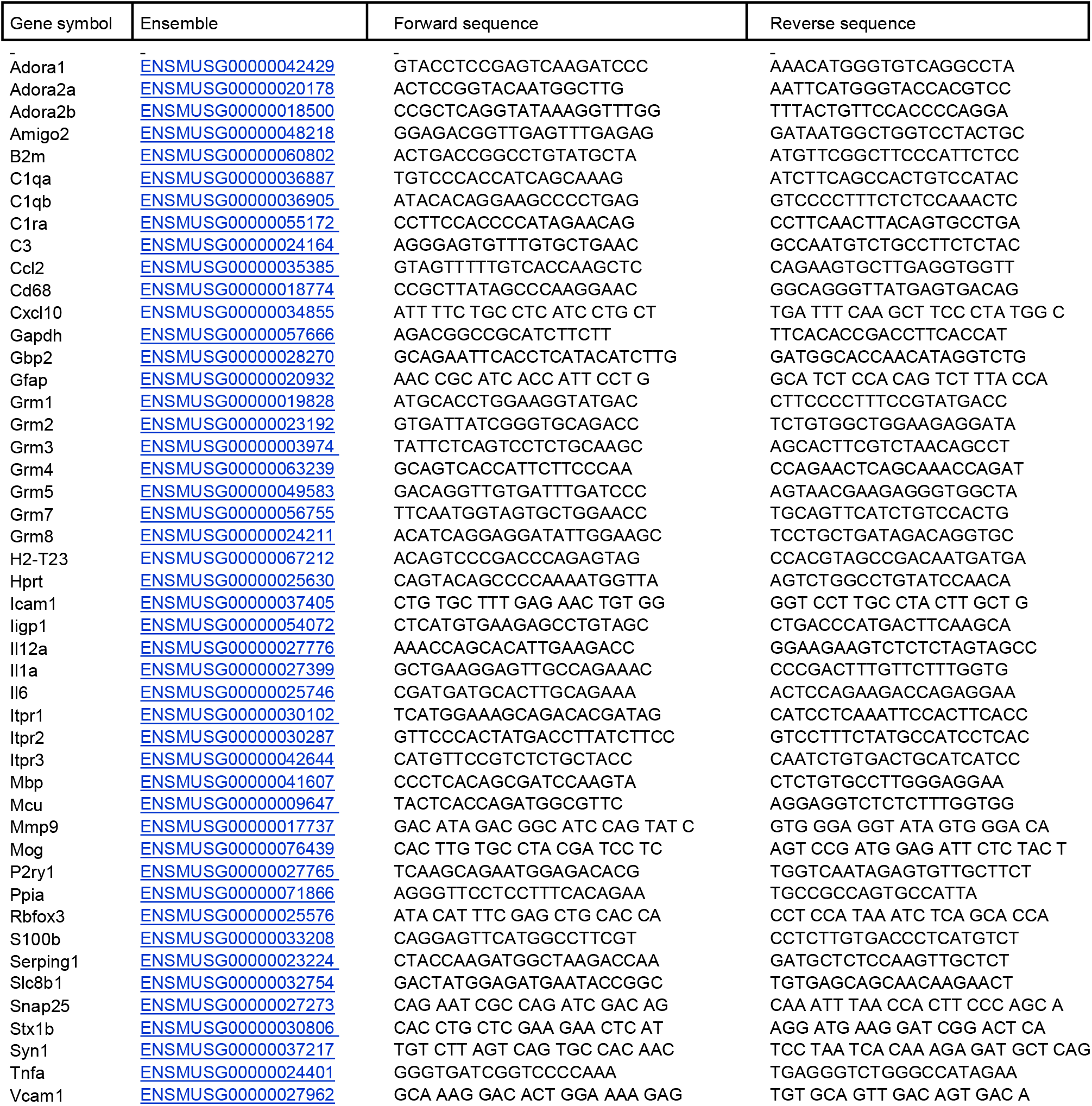
List of primers used for RT-qPCR analysis.

## Bibliography

Akerboom, J., Carreras Calderón, N., Tian, L., Wabnig, S., Prigge, M., Tolö, J., . . . Looger, L. L. (2013). Genetically encoded calcium indicators for multi-color neural activity imaging and combination with optogenetics. Front Mol Neurosci, 6, 2. https://doi.org/10.3389/fnmol.2013.00002

Araque, A., Carmignoto, G., Haydon, P. G., Oliet, S. H., Robitaille, R., & Volterra, A. (2014). Gliotransmitters travel in time and space. Neuron, 81(4), 728–739. https://doi.org/10.1016/j.neuron.2014.02.007

Baraibar, A. M., Belisle, L., Marsicano, G., Matute, C., Mato, S., Araque, A., & Kofuji, P. (2022). Spatial organization of neuron-astrocyte interactions in the somatosensory cortex. Cereb Cortex. https://doi.org/10.1093/cercor/bhac357

Bojarskaite, L., Bjørnstad, D. M., Pettersen, K. H., Cunen, C., Hermansen, G. H., Åbjørsbråten, K. S., . . . Nagelhus, E. A. (2020). Astrocytic Ca^2+^ signaling is reduced during sleep and is involved in the regulation of slow wave sleep. Nat Commun, 11(1), 3240. https://doi.org/10.1038/s41467-020-17062-2

Bosson, A., Boisseau, S., Buisson, A., Savasta, M., Albrieux, M. (2015) Disruption of dopaminergic transmission remodels tripartite synapse morphology and astrocytic calcium activity within substantia nigra pars reticulata. Glia, 63(4):673–83. https://doi.org/10.1002/glia.22777

Calabrese, M., Magliozzi, R., Ciccarelli, O., Geurts, J. J., Reynolds, R., & Martin, R. (2015). Exploring the origins of grey matter damage in multiple sclerosis. Nat Rev Neurosci, 16(3), 147–158. https://doi.org/10.1038/nrn3900

Calabrese, M., Rocca, M. A., Atzori, M., Mattisi, I., Favaretto, A., Perini, P., . . . Filippi, M. (2010). A 3-year magnetic resonance imaging study of cortical lesions in relapse-onset multiple sclerosis. Ann Neurol, 67(3), 376–383. https://doi.org/10.1002/ana.21906

Centonze, D., Bari, M., Rossi, S., Prosperetti, C., Furlan, R., Fezza, F., . . . Maccarrone, M. (2007). The endocannabinoid system is dysregulated in multiple sclerosis and in experimental autoimmune encephalomyelitis. Brain, 130(Pt 10), 2543–2553. https://doi.org/10.1093/brain/awm160

Centonze, D., Muzio, L., Rossi, S., Cavasinni, F., De Chiara, V., Bergami, A., . . . Martino, G. (2009). Inflammation triggers synaptic alteration and degeneration in experimental autoimmune encephalomyelitis. J Neurosci, 29(11), 3442–3452. https://doi.org/10.1523/JNEUROSCI.5804-08.2009

Chao, C. C., Gutiérrez-Vázquez, C., Rothhammer, V., Mayo, L., Wheeler, M. A., Tjon, E. C., . . . Quintana, F. J. (2019). Metabolic control of astrocyte pathogenic activity via cPLA2-MAVS. Cell, 179(7), 1483–1498.e1422. https://doi.org/10.1016/j.cell.2019.11.016

Clarke, L. E., Liddelow, S. A., Chakraborty, C., Münch, A. E., Heiman, M., & Barres, B. A. (2018). Normal aging induces A1-like astrocyte reactivity. Proc Natl Acad Sci U S A, 115(8), E1896–E1905. https://doi.org/10.1073/pnas.1800165115

De Stefano, N., Matthews, P. M., Filippi, M., Agosta, F., De Luca, M., Bartolozzi, M. L., . . . Smith, S. M. (2003). Evidence of early cortical atrophy in MS: relevance to white matter changes and disability. Neurology, 60(7), 1157–1162. https://doi.org/10.1212/01.wnl.0000055926.69643.03

Delekate, A., Füchtemeier, M., Schumacher, T., Ulbrich, C., Foddis, M., & Petzold, G. C. (2014). Metabotropic P2Y1 receptor signalling mediates astrocytic hyperactivity in vivo in an Alzheimer’s disease mouse model. Nat Commun, 5, 5422. https://doi.org/10.1038/ncomms6422

Di Castro, M. A., Chuquet, J., Liaudet, N., Bhaukaurally, K., Santello, M., Bouvier, D., . . . Volterra, A. (2011). Local Ca^2+^ detection and modulation of synaptic release by astrocytes. Nat Neurosci, 14(10), 1276–1284. https://doi.org/10.1038/nn.2929

Ding, F., O’Donnell, J., Thrane, A. S., Zeppenfeld, D., Kang, H., Xie, L., . . . Nedergaard, M. (2013). α1-Adrenergic receptors mediate coordinated Ca^2+^ signaling of cortical astrocytes in awake, behaving mice. Cell Calcium, 54(6), 387–394. https://doi.org/10.1016/j.ceca.2013.09.001

Ding, S., Fellin, T., Zhu, Y., Lee, S. Y., Auberson, Y. P., Meaney, D. F., . . . Haydon, P. G. (2007). Enhanced astrocytic Ca^2+^ signals contribute to neuronal excitotoxicity after status epilepticus. J Neurosci, 27(40), 10674–10684. https://doi.org/10.1523/JNEUROSCI.2001-07.2007

Durkee, C. A., Covelo, A., Lines, J., Kofuji, P., Aguilar, J., & Araque, A. (2019). G_i/o_ protein-coupled receptors inhibit neurons but activate astrocytes and stimulate gliotransmission. Glia, 67(6), 1076–1093. https://doi.org/10.1002/glia.23589

Eljaschewitsch, E., Witting, A., Mawrin, C., Lee, T., Schmidt, P. M., Wolf, S., . . . Ullrich, O. (2006). The endocannabinoid anandamide protects neurons during CNS inflammation by induction of MKP-1 in microglial cells. Neuron, 49(1), 67–79. https://doi.org/10.1016/j.neuron.2005.11.027

Ellwardt, E., Pramanik, G., Luchtman, D., Novkovic, T., Jubal, E. R., Vogt, J., . . . Stroh, A. (2018). Maladaptive cortical hyperactivity upon recovery from experimental autoimmune encephalomyelitis. Nat Neurosci, 21(10), 1392–1403. https://doi.org/10.1038/s41593-018-0193-2

Escartin, C., Galea, E., Lakatos, A., O’Callaghan, J. P., Petzold, G. C., Serrano-Pozo, A., . . . Verkhratsky, A. (2021). Reactive astrocyte nomenclature, definitions, and future directions. Nat Neurosci, 24(3), 312–325. https://doi.org/10.1038/s41593-020-00783-4

Eshaghi, A., Prados, F., Brownlee, W. J., Altmann, D. R., Tur, C., Cardoso, M. J., . . . group, M. s. (2018). Deep gray matter volume loss drives disability worsening in multiple sclerosis. Ann Neurol, 83(2), 210–222. https://doi.org/10.1002/ana.25145

Feinstein, A., Magalhaes, S., Richard, J. F., Audet, B., & Moore, C. (2014). The link between multiple sclerosis and depression. Nat Rev Neurol, 10(9), 507–517. https://doi.org/10.1038/nrneurol.2014.139

Fellin, T., Pascual, O., Gobbo, S., Pozzan, T., Haydon, P. G., & Carmignoto, G. (2004). Neuronal synchrony mediated by astrocytic glutamate through activation of extrasynaptic NMDA receptors. Neuron, 43(5), 729–743. https://doi.org/10.1016/j.neuron.2004.08.011

Habbas, S., Santello, M., Becker, D., Stubbe, H., Zappia, G., Liaudet, N., . . . Volterra, A. (2015). Neuroinflammatory TNFα Impairs Memory via Astrocyte Signaling. Cell, 163(7), 1730–1741. https://doi.org/10.1016/j.cell.2015.11.023

Hamby, M. E., Coppola, G., Ao, Y., Geschwind, D. H., Khakh, B. S., & Sofroniew, M. V. (2012). Inflammatory mediators alter the astrocyte transcriptome and calcium signaling elicited by multiple G-protein-coupled receptors. J Neurosci, 32(42), 14489–14510. https://doi.org/10.1523/JNEUROSCI.1256-12.2012

Han, J., Kesner, P., Metna-Laurent, M., Duan, T., Xu, L., Georges, F., . . . Zhang, X. (2012). Acute cannabinoids impair working memory through astroglial CB1 receptor modulation of hippocampal LTD. Cell, 148(5), 1039–1050. https://doi.org/10.1016/j.cell.2012.01.037

Hartmann, K., Sepulveda-Falla, D., Rose, I. V. L., Madore, C., Muth, C., Matschke, J., . . . Krasemann, S. (2019). Complement 3^+^-astrocytes are highly abundant in prion diseases, but their abolishment led to an accelerated disease course and early dysregulation of microglia. Acta Neuropathol Commun, 7(1), 83. https://doi.org/10.1186/s40478-019-0735-1

Hasel, P., Rose, I. V. L., Sadick, J. S., Kim, R. D., & Liddelow, S. A. (2021). Neuroinflammatory astrocyte subtypes in the mouse brain. Nat Neurosci, 24(10), 1475–1487. https://doi.org/10.1038/s41593-021-00905-6

Haustein, M. D., Kracun, S., Lu, X. H., Shih, T., Jackson-Weaver, O., Tong, X., . . . Khakh, B. S. (2014). Conditions and constraints for astrocyte calcium signaling in the hippocampal mossy fiber pathway. Neuron, 82(2), 413–429. https://doi.org/10.1016/j.neuron.2014.02.041

Hirrlinger, P. G., Scheller, A., Braun, C., Hirrlinger, J., & Kirchhoff, F. (2006). Temporal control of gene recombination in astrocytes by transgenic expression of the tamoxifen-inducible DNA recombinase variant CreERT2. Glia, 54(1), 11–20. https://doi.org/10.1002/glia.20342

Hou, B., Zhang, Y., Liang, P., He, Y., Peng, B., Liu, W., . . . He, X. (2020). Inhibition of the NLRP3-inflammasome prevents cognitive deficits in experimental autoimmune encephalomyelitis mice via the alteration of astrocyte phenotype. Cell Death Dis, 11(5), 377. https://doi.org/10.1038/s41419-020-2565-2

Jafari, M., Schumacher, A. M., Snaidero, N., Ullrich Gavilanes, E. M., Neziraj, T., Kocsis-Jutka, V., . . . Kerschensteiner, M. (2021). Phagocyte-mediated synapse removal in cortical neuroinflammation is promoted by local calcium accumulation. Nat Neurosci, 24(3), 355–367. https://doi.org/10.1038/s41593-020-00780-7

Jiang, R., Diaz-Castro, B., Looger, L. L., & Khakh, B. S. (2016). Dysfunctional calcium and glutamate signaling in striatal astrocytes from Huntington’s disease model mice. J Neurosci, 36(12), 3453–3470. https://doi.org/10.1523/JNEUROSCI.3693-15.2016

Jürgens, T., Jafari, M., Kreutzfeldt, M., Bahn, E., Brück, W., Kerschensteiner, M., & Merkler, D. (2016). Reconstruction of single cortical projection neurons reveals primary spine loss in multiple sclerosis. Brain, 139(Pt 1), 39–46. https://doi.org/10.1093/brain/awv353

Kuchibhotla, K. V., Lattarulo, C. R., Hyman, B. T., & Bacskai, B. J. (2009). Synchronous hyperactivity and intercellular calcium waves in astrocytes in Alzheimer mice. Science, 323(5918), 1211–1215. https://doi.org/10.1126/science.1169096

Lagumersindez-Denis, N., Wrzos, C., Mack, M., Winkler, A., van der Meer, F., Reinert, M. C., . . . Nessler, S. (2017). Differential contribution of immune effector mechanisms to cortical demyelination in multiple sclerosis. Acta Neuropathol, 134(1), 15–34. https://doi.org/10.1007/s00401-017-1706-x

Liddelow, S. A., Guttenplan, K. A., Clarke, L. E., Bennett, F. C., Bohlen, C. J., Schirmer, L., . . . Barres, B. A. (2017). Neurotoxic reactive astrocytes are induced by activated microglia. Nature, 541(7638), 481–487. https://doi.org/10.1038/nature21029

Lines, J., Baraibar, A. M., Fang, C., Martin, E. D., Aguilar, J., Lee, M. K., . . . Kofuji, P. (2022). Astrocyte-neuronal network interplay is disrupted in Alzheimer’s disease mice. Glia, 70(2), 368–378. https://doi.org/10.1002/glia.24112

Lines, J., Martin, E. D., Kofuji, P., Aguilar, J., & Araque, A. (2020). Astrocytes modulate sensory-evoked neuronal network activity. Nat Commun, 11(1), 3689. https://doi.org/10.1038/s41467-020-17536-3

Linnerbauer, M., Wheeler, M. A., & Quintana, F. J. (2020). Astrocyte Crosstalk in CNS Inflammation. Neuron, 108(4), 608–622. https://doi.org/10.1016/j.neuron.2020.08.012

Lütcke, H., Murayama, M., Hahn, T., Margolis, D. J., Astori, S., Zum Alten Borgloh, S. M., . . . Hasan, M. T. (2010). Optical recording of neuronal activity with a genetically-encoded calcium indicator in anesthetized and freely moving mice. Front Neural Circuits, 4, 9. https://doi.org/10.3389/fncir.2010.00009

Mahad, D. H., Trapp, B. D., & Lassmann, H. (2015). Pathological mechanisms in progressive multiple sclerosis. Lancet Neurol, 14(2), 183–193. https://doi.org/10.1016/S1474-4422(14)70256-X

Mandolesi, G., Musella, A., Gentile, A., Grasselli, G., Haji, N., Sepman, H., . . . Centonze, D. (2013). Interleukin-1β alters glutamate transmission at purkinje cell synapses in a mouse model of multiple sclerosis. J Neurosci, 33(29), 12105–12121. https://doi.org/10.1523/JNEUROSCI.5369-12.2013

Marsicano, G., Wotjak, C. T., Azad, S. C., Bisogno, T., Rammes, G., Cascio, M. G., . . . Lutz, B. (2002). The endogenous cannabinoid system controls extinction of aversive memories. Nature, 418(6897), 530–534. https://doi.org/10.1038/nature00839

Martínez-Gallego, I., Pérez-Rodríguez, M., Coatl-Cuaya, H., Flores, G., & Rodríguez-Moreno, A. (2022). Adenosine and astrocytes determine the developmental dynamics of spike timing-dependent plasticity in the somatosensory cortex. J Neurosci, 42(31), 6038–6052. https://doi.org/10.1523/JNEUROSCI.0115-22.2022

Min, R., & Nevian, T. (2012). Astrocyte signaling controls spike timing-dependent depression at neocortical synapses. Nat Neurosci, 15(5), 746–753. https://doi.org/10.1038/nn.3075

Moreno-García, Á., Bernal-Chico, A., Colomer, T., Rodríguez-Antigüedad, A., Matute, C., & Mato, S. (2020). Gene expression analysis of astrocyte and microglia endocannabinoid signaling during autoimmune demyelination. Biomolecules, 10(9). https://doi.org/10.3390/biom10091228

Nagai, J., Bellafard, A., Qu, Z., Yu, X., Ollivier, M., Gangwani, M. R., . . . Khakh, B. S. (2021). Specific and behaviorally consequential astrocyte G_q_ GPCR signaling attenuation in vivo with iβARK. Neuron, 109(14), 2256–2274.e2259. https://doi.org/10.1016/j.neuron.2021.05.023

Nanclares, C., Poynter, J., Martell-Martinez, H. A., Vermilyea, S., Araque, A., Kofuji, P., . . . Covelo, A. (2023). Dysregulation of astrocytic Ca^2+^ signaling and gliotransmitter release in mouse models of α-synucleinopathies. Acta Neuropathol. https://doi.org/10.1007/s00401-023-02547-3

Navarrete, M., & Araque, A. (2010). Endocannabinoids potentiate synaptic transmission through stimulation of astrocytes. Neuron, 68(1), 113–126. https://doi.org/10.1016/j.neuron.2010.08.043

Ohkura, M., Sasaki, T., Sadakari, J., Gengyo-Ando, K., Kagawa-Nagamura, Y., Kobayashi, C., . . . Nakai, J. (2012). Genetically encoded green fluorescent Ca2+ indicators with improved detectability for neuronal Ca^2+^ signals. PLoS One, 7(12), e51286. https://doi.org/10.1371/journal.pone.0051286

Perea, G., & Araque, A. (2005). Properties of synaptically evoked astrocyte calcium signal reveal synaptic information processing by astrocytes. J Neurosci, 25(9), 2192–2203. https://doi.org/10.1523/JNEUROSCI.3965-04.2005

Pirttimaki, T. M., Sims, R. E., Saunders, G., Antonio, S. A., Codadu, N. K., & Parri, H. R. (2017). Astrocyte-mediated neuronal synchronization properties revealed by false gliotransmitter release. J Neurosci, 37(41), 9859–9870. https://doi.org/10.1523/JNEUROSCI.2761-16.2017.

Pitt, D., Werner, P., & Raine, C. S. (2000). Glutamate excitotoxicity in a model of multiple sclerosis. Nat Med, 6(1), 67–70. https://doi.org/10.1038/71555

Ponath, G., Park, C., & Pitt, D. (2018). The role of astrocytes in multiple sclerosis. Front Immunol, 9, 217. https://doi.org/10.3389/fimmu.2018.00217

Potter, L. E., Paylor, J. W., Suh, J. S., Tenorio, G., Caliaperumal, J., Colbourne, F., . . . Kerr, B. J. (2016). Altered excitatory-inhibitory balance within somatosensory cortex is associated with enhanced plasticity and pain sensitivity in a mouse model of multiple sclerosis. J Neuroinflammation, 13(1), 142. https://doi.org/10.1186/s12974-016-0609-4

Qin, H., He, W., Yang, C., Li, J., Jian, T., Liang, S., . . . Zhang, K. (2020). onitoring astrocytic Ca^2+^ activity in freely behaving mice. Front Cell Neurosci, 14, 603095. https://doi.org/10.3389/fncel.2020.603095

Robin, L. M., Oliveira da Cruz, J. F., Langlais, V. C., Martin-Fernandez, M., Metna-Laurent, M., Busquets-Garcia, A., . . . Marsicano, G. (2018). Astroglial CB_1_ receptors determine synaptic D-serine availability to enable recognition memory. Neuron, 98(5), 935–944.e935. https://doi.org/10.1016/j.neuron.2018.04.034

Rothstein, J. D., Martin, L., Levey, A. I., Dykes-Hoberg, M., Jin, L., Wu, D., . . . Kuncl, R. W. (1994). Localization of neuronal and glial glutamate transporters. Neuron, 13(3), 713–725. https://doi.org/10.1016/0896-6273(94)90038-8

Serrat, R., Covelo, A., Kouskoff, V., Delcasso, S., Ruiz-Calvo, A., Chenouard, N., . . . Marsicano, G. (2021). Astroglial ER-mitochondria calcium transfer mediates endocannabinoid-dependent synaptic integration. Cell Rep, 37(12), 110133. https://doi.org/10.1016/j.celrep.2021.110133

Shah, D., Gsell, W., Wahis, J., Luckett, E. S., Jamoulle, T., Vermaercke, B., . . . De Strooper, B. (2022). Astrocyte calcium dysfunction causes early network hyperactivity in Alzheimer’s disease. Cell Rep, 40(8), 111280. https://doi.org/10.1016/j.celrep.2022.111280

Shigetomi, E., Patel, S., & Khakh, B. S. (2016). Probing the complexities of astrocyte calcium signaling. Trends Cell Biol, 26(4), 300–312. https://doi.org/10.1016/j.tcb.2016.01.003

Sofroniew, M. V. (2020). strocyte Reactivity: Subtypes, states, and functions in CNS innate immunity. Trends Immunol, 41(9), 758–770. https://doi.org/10.1016/j.it.2020.07.004

Stobart, J. L., Ferrari, K. D., Barrett, M. J. P., Stobart, M. J., Looser, Z. J., Saab, A. S., & Weber, B. (2018). Long-term in vivo calcium imaging of astrocytes reveals distinct cellular compartment responses to sensory stimulation. Cereb Cortex, 28(1), 184–198. https://doi.org/10.1093/cercor/bhw366

Sun, W., McConnell, E., Pare, J. F., Xu, Q., Chen, M., Peng, W., . . . Nedergaard, M. (2013). Glutamate-dependent neuroglial calcium signaling differs between young and adult brain. Science, 339(6116), 197–200. https://doi.org/10.1126/science.1226740

Talantova, M., Sanz-Blasco, S., Zhang, X., Xia, P., Akhtar, M. W., Okamoto, S., . . . Lipton, S. A. (2013). Aβ induces astrocytic glutamate release, extrasynaptic NMDA receptor activation, and synaptic loss. Proc Natl Acad Sci U S A, 110(27), E2518–2527. https://doi.org/10.1073/pnas.1306832110

Wang, X., Lou, N., Xu, Q., Tian, G. F., Peng, W. G., Han, X., . . . Nedergaard, M. (2006). Astrocytic Ca^2+^ signaling evoked by sensory stimulation in vivo. Nat Neurosci, 9(6), 816–823. https://doi.org/10.1038/nn1703

Wujek, J. R., Bjartmar, C., Richer, E., Ransohoff, R. M., Yu, M., Tuohy, V. K., & Trapp, B. D. (2002). Axon loss in the spinal cord determines permanent neurological disability in an animal model of multiple sclerosis. J Neuropathol Exp Neurol, 61(1), 23–32. https://doi.org/10.1093/jnen/61.1.23

Yang, G., Parkhurst, C. N., Hayes, S., & Gan, W. B. (2013). Peripheral elevation of TNF-α leads to early synaptic abnormalities in the mouse somatosensory cortex in experimental autoimmune encephalomyelitis. Proc Natl Acad Sci U S A, 110(25), 10306–10311. https://doi.org/10.1073/pnas.1222895110

Zur Nieden, R., & Deitmer, J. W. (2006). The role of metabotropic glutamate receptors for the generation of calcium oscillations in rat hippocampal astrocytes in situ. Cereb Cortex, 16(5), 676–687. https://doi.org/10.1093/cercor/bhj013

